# Mapping *Plasmodium* transitions and interactions in the *Anopheles* female

**DOI:** 10.1101/2024.11.12.623125

**Authors:** Yan Yan, Lisa H. Verzier, Elaine Cheung, Federico Appetecchia, Sandra March, Ailsa R. Craven, Esrah Du, Alexandra S. Probst, Tasneem A. Rinvee, Laura E. de Vries, Jamie Kauffman, Sangeeta N. Bhatia, Elisabeth Nelson, Naresh Singh, Duo Peng, W. Robert Shaw, Flaminia Catteruccia

## Abstract

The human malaria parasite, *Plasmodium falciparum*, relies exclusively on *Anopheles* mosquitoes for transmission. Once ingested during blood feeding, most parasites die in the mosquito midgut lumen or during epithelium traversal^1^. How surviving ookinetes interact with midgut cells and form oocysts is poorly known, yet these steps are essential to initiate a remarkable growth process culminating in the production of thousands of infectious sporozoites^2^. Here, using single-cell RNA sequencing of both parasites and mosquito cells across different developmental stages and metabolic conditions, we unveil key transitions and mosquito-parasite interactions occurring in the midgut. Functional analyses uncover processes regulating oocyst growth and identify the transcription factor *Pf*SIP2 as essential for sporozoite infection of human hepatocytes. Combining shared mosquito-parasite barcode analysis with confocal microscopy, we reveal that parasites preferentially interact with midgut progenitor cells during epithelial crossing, potentially using their basal location as an exit landmark. Additionally, we show tight connections between extracellular late oocysts and surrounding muscle cells that may ensure parasites adherence to the midgut. We confirm our major findings in several mosquito-parasite combinations, including field-derived parasites. Our study provides fundamental insight into the molecular events characterizing previously inaccessible biological transitions and mosquito-parasite interactions, and identifies candidates for transmission-blocking strategies.

## Introduction

The apicomplexan parasite *Plasmodium falciparum* is responsible for 90% of cases of malaria, a disease that in 2023 alone caused the death of close to 600,000 people, mostly young children in sub-Saharan Africa^3^. Like all malaria parasites, *P. falciparum* requires a blood-feeding vector for transmission between humans, and over the course of a long co-evolutionary history has adapted to be exclusively transmitted by mosquitoes of the *Anopheles* genus. The parasite journey in the *Anopheles* female starts when male and female gametocytes are ingested during a blood meal and rapidly mature into gametes, leading to fertilization and formation of a zygote which transforms into a motile ookinete in the blood bolus. Between 24–36 hours after ingestion, ookinetes must cross the midgut epithelium to avoid killing during blood digestion. These initial steps result in substantial parasite loss, representing one of the most severe bottlenecks in the parasite lifecycle^1^. After reaching the basal side of the midgut, ookinetes round up via an intermediate took stage (‘**t**ransition from **ook**inete’)^4^ to transform into extracellular oocysts, which over the next 7–10 days will extensively grow in size just underneath the basal lamina surrounding the midgut epithelial layer while undergoing DNA replication. This remarkable growth process culminates in the formation of thousands of infectious sporozoites, an intricate process where individual parasites must be segmented precisely over a relatively short period of time^2^ (Fig. 1a). When mature, sporozoites egress from oocysts and invade the mosquito salivary glands, from where they can be transmitted to the next person when the mosquito bites again.

**Fig. 1.**
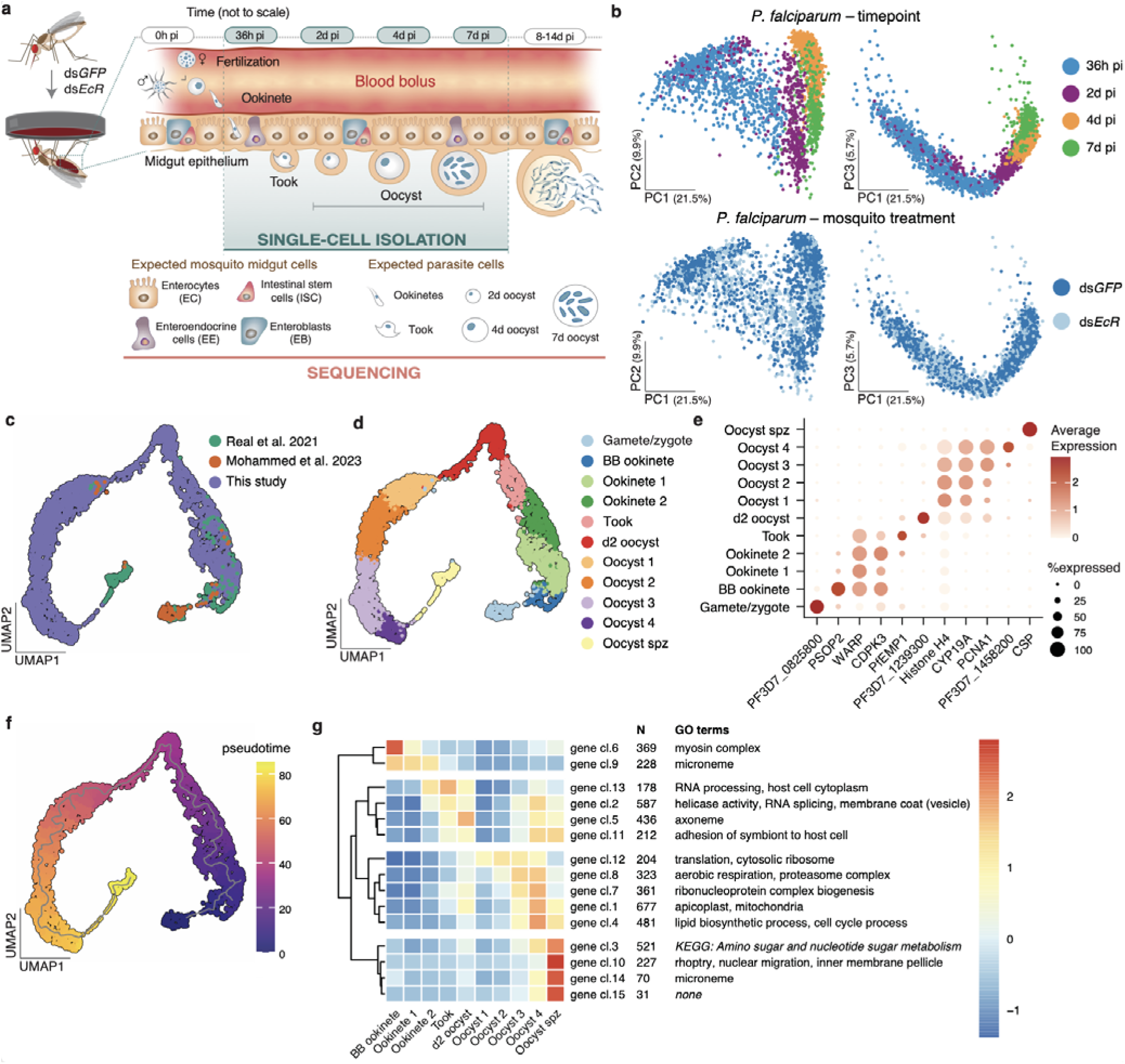
Single cell RNA-seq provides a complete map of *P. falciparum* midgut stages. **a,** Schematic of dual single-cell RNA sequencing (dual scRNA-seq) displaying expected parasite forms and mosquito midgut cell types. Single *P. falciparum* parasites and *An. gambiae* midgut cells were collected from GFP- (control) and EcR-depleted mosquitoes at the time points indicated by the shaded area: 36h post infection (h pi), 2 days post infection (d pi), 4d pi, and 7d pi. and across four biological replicates **b,** PCA plots of the 3,495 parasites collected across the four time points (top) and the two treatments (bottom). **c-d,** UMAP plots (**c**) integrating our scRNA-seq data with midgut stage data from two other studies^21,22^, and (**d**) showing 11 distinct parasite clusters. **e,** Dot plot of the top marker genes for each of the parasite clusters, showing their normalized average expression (color) and the proportion of parasites expressing each gene (size). **f,** Pseudotime analysis of the integrated scRNA-seq data with trajectory (grey line). **g,** Gene cluster analysis of our expression data identifies four main branches of co-expressed genes across the different clusters (gene cl: gene cluster, N: number of genes in the cluster). Selected GO terms and KEGG annotations enriched in each gene cluster are indicated whenever present. Mosquito schematics in **a** are adapted from ref. 36 under a Creative Commons licence CC BY 4.0 and parasites were reprinted from ref 24 with permission from Elsevier.

The mosquito stages of *P. falciparum* development are relatively poorly characterized compared to asexual blood stages, despite their essential role in transmission, and current knowledge is largely extrapolated from work on more tractable rodent malaria models^5–10^. While a wealth of studies have visualized the route by which ookinetes invade the midgut epithelium^11–15^, whether they interact with specific midgut cell types and the mechanisms that govern their transition into oocysts remain largely unknown. The processes fueling oocyst development and promoting sporozoite formation are also not fully understood. Tackling these questions has been challenging, as mosquito stages of *P. falciparum* are restricted to *in vivo* study and have limited genetic tools relative to asexual blood stage parasites^16^. Moreover, the low ookinete and oocyst load in a midgut limits genome-wide multi-omics approaches of those stages, which have been therefore mostly confined to non-human *Plasmodium* species where parasite densities are substantially higher^17–20^. For similar reasons, single-cell RNA sequencing (scRNA-seq) studies of *P. falciparum* in the mosquito have focused on stages where parasites are numerous and readily isolated (i.e. blood bolus stages and sporozoites)^21,22^, leaving significant gaps in the parasite life cycle.

Here, we reconstruct critical bottleneck stages of parasite growth in the midgut and unveil novel mosquito-parasite interactions by performing dual scRNA-seq of parasites and midgut cells over time, capturing key processes mediating midgut crossing, oocyst growth, and the initial stages of sporozoite formation.

## Results

### ScRNA-seq of midgut and parasite cells

We infected *Anopheles gambiae* – the major malaria vector in sub-Saharan Africa – with *P. falciparum* and isolated both parasites and midgut cells at 4 different time points: invading ookinetes (36 hours post infection, h pi); newly formed oocysts (2 days pi, d pi); growing oocysts (4d pi) and late oocysts that may have begun sporozoite segmentation (7d pi) (Fig. 1a; Extended Data Fig. 1a). We did not attempt to isolate oocysts at later time points given their large size (>50 µm) exceeds the limits of 10X technology. Due to low parasite numbers relative to the overwhelming number of mosquito cells^23^, we enriched for parasites by optimizing the single-cell isolation protocol (detailed in the methods) and applied deeper sequencing. Briefly, infected midguts were partially digested with collagenase IV and elastase, then filtered through a series of cell strainers to remove large mosquito cell clumps. The resulting single-cell suspensions contained both midgut cells and parasites, with cell viability over 93% as determined by trypan blue staining. We compared parasites developing in mosquitoes under two metabolic conditions: control mosquitoes (injected with double-stranded RNA against *eGFP*, ds*GFP*) and mosquitoes depleted of the Ecdysone Receptor (ds*EcR*), the nuclear co-receptor of the steroid hormone 20-hydroxy-ecdysone (20E)^24^ (Extended Data Fig. 1b-e). Impairing 20E signaling reliably accelerated oocyst growth, which we reasoned would provide more granular resolution at the later stages, and did not affect parasites numbers as previously observed at low parasite densities^24,25^ (Extended Data Fig. 1b, c). After removing low-quality parasites with low transcript and gene counts and high mitochondrial percentage, we successfully profiled 3,495 parasites, detecting a median of 242–1,773 genes at the various time points and treatments (Extended Data Fig. 1f, Supplementary Table 1).

Principal component analysis (PCA) revealed a large separation along the first component, with parasites at 2, 4 and 7d pi clustering away from 36h pi parasites (Fig. 1b). The ookinete-oocyst transition (2d vs. 36h) was characterized by a downregulation of RNA metabolism paralleled by an upregulation of translation, while oocyst growth was marked by increased aerobic respiration (4d vs. 2d) (Extended Data Fig. 2, Supplementary Table 2). PCA at early time points showed no clear separation between parasites developing in ds*GFP* and ds*EcR* mosquitoes, and consistently, there was no difference in gene expression between these groups. At 7d pi, however, we detected the upregulation of several genes in parasites from ds*EcR* females (Extended Data Fig. 2, Supplementary Table 2), likely reflecting the more advanced oocyst stage in this group (Extended Data Fig. 1b). Among upregulated genes were the circumsporozoite protein (CSP, the most abundant sporozoite surface protein), and genes from the rhoptries, secretory organelles essential for invasion of host cells^26^, suggesting we successfully captured the start of sporozoite segmentation.

### Parasite atlas across all midgut stages

Next, we generated a Uniform Manifold Approximation and Projection (UMAP) of parasites in the mosquito midgut by integrating our datasets with parasites from previous scRNA-seq studies^21,22^, including gametes and blood bolus (BB) ookinetes as well as sporozoites isolated from oocysts (Fig. 1c). This complete midgut atlas uncovered 11 parasite clusters, which we annotated based on marker genes and data source as follows: Gametes/Zygotes, BB ookinetes, invading ookinetes (Ookinete 1 and 2), the ookinete-oocyst transition stage (Took), newly formed oocysts (d2 oocyst), growing oocysts (Oocyst 1-4) and oocysts that are segmenting into sporozoites (Oocyst spz) (Fig. 1d, e, Extended Data Fig. 3a, Supplementary Table 3). In agreement with our observation of early segmentation signals at 7d pi, a few cells bridged the gap between oocysts in an advanced growth phase and sporozoites isolated from oocysts (Fig. 1c, d, Extended Fig. 3b). Pseudotime analysis revealed a single trajectory aligning with the expected clusters (Fig. 1f).

### Gene clusters unveil key transitions

We clustered genes with similar expression patterns over pseudotime, using only our data due to differences in gene detection rates among scRNA-seq techniques. The 15 gene clusters identified in this analysis separated into four major branches (Fig. 1g, Supplementary Table 4). The first branch (gene clusters 6, 9) captured expected differences between BB ookinetes and ookinetes that are crossing the epithelium. Cluster 6, highly expressed in BB parasites, was associated with ookinete maturation and characterized by genes involved in cytoskeleton and myosin complex^27^. Cluster 9, expressed across all ookinete clusters, comprised micronemal proteins critical for motility and midgut traversal, such as circumsporozoite-and TRAP-related protein (CTRP), chitinase, and glycosylphosphatidylinositol-anchored micronemal antigen (GAMA)^28,29^.

The second branch was largely characterized by processes related to transcription, surface remodeling, and adhesion, which is interesting given this branch encompasses a transition (ookinete-oocyst) during which parasites undergo profound morphological changes. At the onset of this transition (clusters 13 and 2) we detected strong signals of genes encoding for helicase activity and RNA processing, including splicing. Clusters 2 and 5 instead comprised membrane and vesicle trafficking genes which may facilitate protein transport and surface remodeling. Additionally, two clusters in this branch (13 and 11) contained several *var* genes encoding variants of *Pf*EMP1 proteins, well characterized for their adhesion properties in the asexual blood stage^30^. It is intriguing to speculate that young oocysts may use *Pf*EMP1 to anchor themselves between midgut cells and basal lamina, in a manner similar to how asexual parasites adhere to the lining of blood vessels^30^.

The largest branch, spanning from d2 oocysts to Oocyst spz, captured the processes fueling the growth phase, including genes regulating translation, proteasome activity, and ribonucleoprotein complex (clusters 12, 8 and 7) as well as genes involved in aerobic respiration, lipid biosynthesis, and apicoplast functions (clusters 8, 1 and 4). Gene cluster 4, highest in Oocysts 4, included functions related to the cell cycle, like DNA replication (Fig. 1g, Supplementary Table 4).

The final branch, highly expressed in Oocyst spz, was characterized by clear signatures of sporozoite segmentation. Clusters 3, 10, 14 and 15 included genes involved in nuclear migration, components of the inner membrane complex (a scaffolding compartment essential for daughter cell formation) and the cytoskeleton network supporting it, and genes from the invasive organelles rhoptries and micronemes, likely preparing sporozoites for salivary gland and hepatocyte invasion^26,31,32^ (Fig. 1g, Supplementary Table 4).

Although poly-A capture is not designed to target ribosomal RNAs (rRNAs), these molecules are ubiquitously present in *Plasmodium* bulk and single cell RNA results^33^. We detected limited expression of different rRNAs via random binding, organized in distinct gene expression patterns (Extended Data Fig. 3c). Expression of the A1/A2 forms that are dominant in asexual parasites^34^ appeared highest in ookinete clusters, then gradually decreased—though not entirely to zero—in later stages. The two mosquito-stage rRNAs, S1 and S2^34^, were expressed throughout oocyst growth, with S1 starting in Took and S2 in d2 oocyst.

Combined, these gene expression networks provide significant information on the key developmental transitions that parasites undergo in the mosquito midgut, shedding light into the biology of these understudied stages.

### *Pf*ATP4 and *Pf*LRS regulate oocyst growth

To assess the functional role of factors emerging from the gene network analysis, we prioritized candidates from different gene clusters that have known potent inhibitors. We selected *Pf*ATP4, an Na^+^-ATPase essential for sodium ion efflux (gene cluster 2) which is inhibited by the drug cipargamin (CIP, original name: NITD609), and the leucine-tRNA ligase (*Pf*LRS, gene cluster 1), which is inhibited by the compound MMV670325 (original name: AN6426) (Fig. 1g, Fig. 2a). We provided these drugs to infected females via sugar solution, from the onset of oocyst formation (2d pi) until 10d pi (Fig. 2b). While neither drug reduced oocyst intensity or prevalence (Extended Data Fig. 4a), both CIP and MMV670325 considerably impaired oocyst growth. As a result, we detected no sporozoites in the salivary glands of CIP-treated mosquitoes at 14d pi and observed a 54% reduction in sporozoite prevalence after MMV670325 ingestion relative to controls (Fig. 2b). Shorter exposure times of CIP (2-4d pi and 2-7d pi) gave similar results, suggesting that regulation of sodium efflux is essential during the early phases of oocyst growth, and the effects were dose-dependent (Fig. 2c, d, Extended Data Fig. 4b-d).

**Fig. 2.**
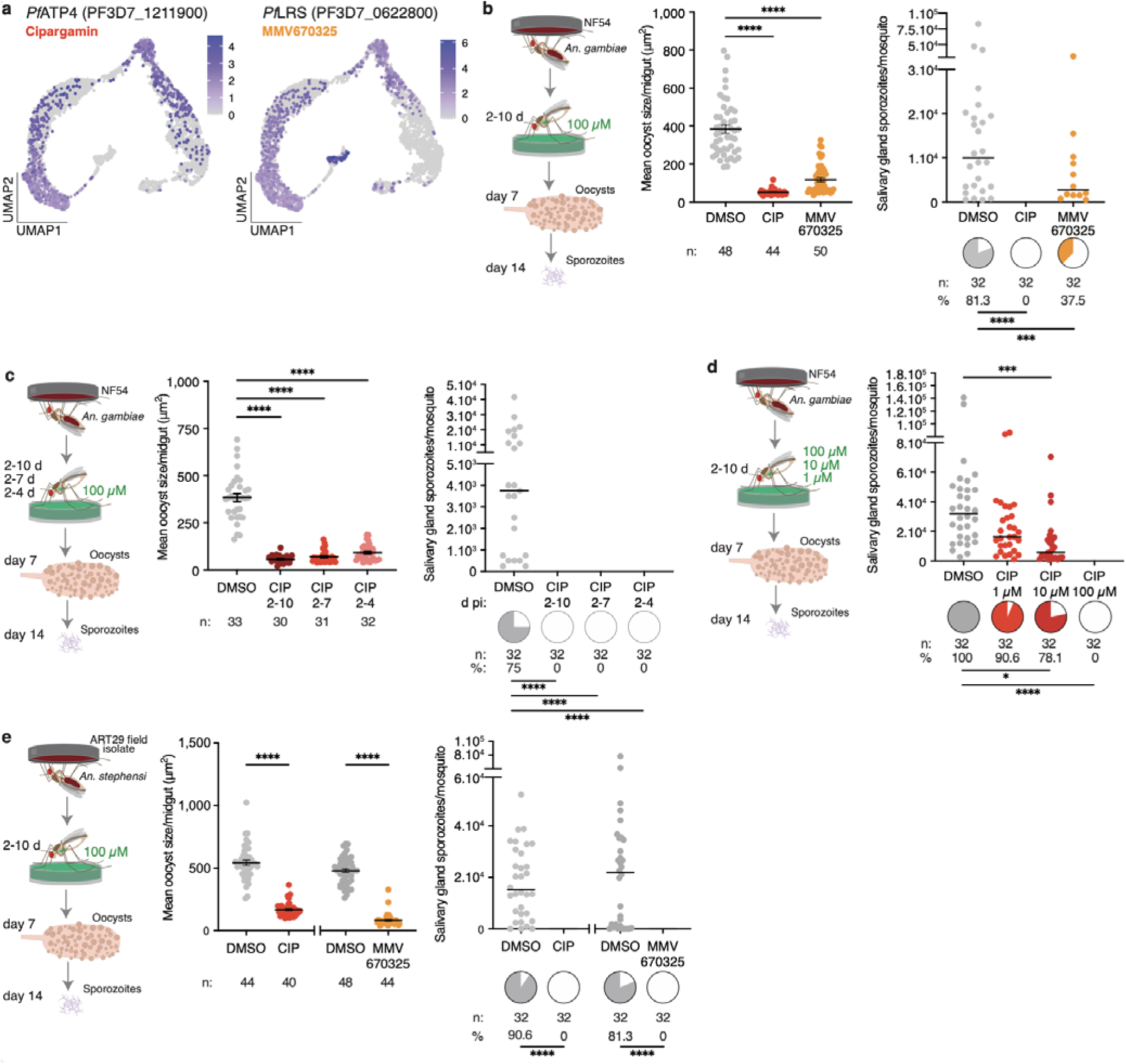
Functional analysis of candidate genes. **a,** Expression profile of *Pf*ATP4, target of cipargamin (CIP), and *Pf*LRS, target of MMV670325. **b,** *An. gambiae* infected with NF54 parasites were exposed to CIP and MMV670325 via sugar solution at 100 µM delivered from 2d to 10d pi (left panel). Drug ingestion reduced oocyst size at 7d pi (middle panel, p<0.0001) subsequently leading to reduced sporozoite prevalence (pie charts, right panel, p<0.0001 and p=0.008) in salivary glands at 14d pi. **c,** Shortening CIP exposure to 2-7 and 2-4d pi also reduced oocyst size at 7d pi (left and middle panels, p<0.0001), subsequently leading to no sporozoites in the salivary glands by 14d pi (right panel, p<0.0001). **d**, Reducing CIP dosage from 100 µM to 10 and 1 µM in *An. gambiae* infected with NF54 parasites led to a dose-dependent reduction in sporozoite prevalence (p=0.0108 and p<0.0001) and intensity (p<0.0001). **e**, Exposure to CIP and MMV670325 in *An. stephensi* mosquitoes infected with ART29 field isolate (left panel) resulted in comparable reduction in oocyst size at 7d pi (middle panel, p<0.0001) and no sporozoites in the salivary glands at 14d pi (pie charts, right panel, p<0.0001). Number of mosquitoes dissected (*n*) and prevalence of infection (%) are indicated under each sample. Oocyst size data are represented as mean ± SEM, sporozoite intensity as median in panels and both are compared using Kruskal-Wallis test and Dunn’s correction (panels b, c, and d) or Mann Whitney two-tailed (panel e). Infection prevalences represented by pie charts are compared with Fisher’s exact test, two-tailed. Data are pooled from two independent infections. *p<0.05, **p<0.01, ***p<0.001, ****p<0.0001. Schematics in **b**, **c**, **d**, **e** are adapted from ref. 36 under a Creative Commons licence CC BY 4.0.

Similar results were obtained with both drugs in a different mosquito-parasite combination (*Anopheles stephensi* infected with *P. falciparum* ART29, an artemisinin-resistant field isolate from Cambodia), and also when exposing *An. gambiae* females to CIP after infection with *P. falciparum* P5, a polyclonal field isolate from Burkina Faso^25^ (Fig. 2e, Extended Data Fig. 4e, f). These results validate *Pf*ATP4 and *Pf*LRS as essential for oocyst development in *P. falciparum*.

Two additional targets – the elongation factor 2 *Pf*eEF2 (cluster 12) and the lactate/H^+^ transporter *Pf*FNT (cluster 7) – showed smaller effects when mosquitoes were exposed to sugar solutions of their respective inhibitors, MMV643121 and MMV007839 (Fig. 1g, Extended Data Fig. 4g, h). MMV643121 slightly decreased oocyst size at 7d pi but did not affect sporozoite intensity or prevalence, while MMV007839 had no effect on oocyst size but significantly reduced sporozoite intensity, indicating a possible role for this transporter in sporozoite segmentation or salivary gland invasion (Extended Data Fig. 4g, h).

These results highlight how our scRNA-seq data can be systematically mined to uncover essential targets for parasite development. The genes validated here could be targeted in novel mosquito-based malaria control strategies, as previously demonstrated using cytochrome b inhibitors^35,36^.

### *Pf*SIP2 ensures hepatocyte invasion

In the final branch, along with rhoptry and cytoskeletal genes, we also detected the ApiAP2 transcription factor *Pf*SIP2 (gene cluster 10) (Fig. 1g, Fig. 3a). In asexual blood stages, *Pf*SIP2 has an essential role likely in daughter merozoite formation^37^, yet its role during mosquito stages remains unknown. To test its function, we generated a conditional *Pf*SIP2 knockdown line integrating the TetR-DOZI aptamer system at the C-terminus (Extended Data Fig. 5a, b). Removal of anhydrotetracycline (ATc) in asexual blood stage cultures led to parasite death, confirming the essentiality of *Pf*SIP2 (Extended Data Fig. 5c)^37^. We adapted this system to mosquito stages by providing ATc throughout gametocyte cultures and in the mosquito daily sugar solutions, and induced gene knockdown by withdrawing the compound immediately before mosquito infection (Fig. 3b). *Pf*SIP2 parasites were less infectious to mosquitoes than wild type co-cultured ones, but this is often observed in mosquito infections with transgenic parasites that have undergone bottlenecks during the selection process. When we compared *Pf*SIP2 +ATc and –ATc groups, we found no differences in terms of oocyst intensity, prevalence and size at 7d pi, nor in sporozoite intensity and prevalence at 14d pi (Extended Data Fig. 5d-f). ATc withdrawal, however, severely impaired the ability of *Pf*SIP2 sporozoites to invade primary human hepatocytes, inducing a remarkable decrease in both invasion efficiency (67% reduction) and the formation of exoerythrocytic forms (EEFs) (96% reduction) (Fig. 3c, d).

**Fig. 3.**
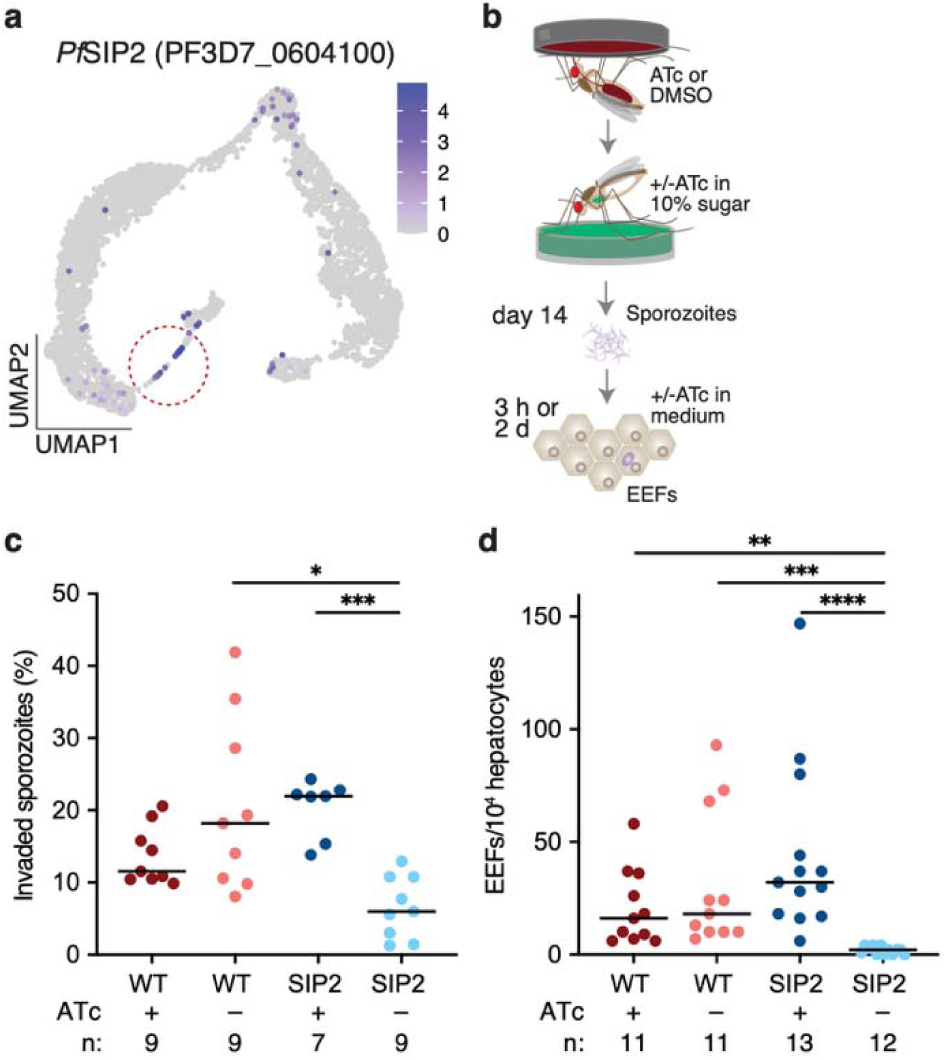
*Pf*SIP2 knockdown impairs hepatocyte infection. **a,** Expression of *Pf*SIP2 (PF3D7_0604100) is highest in segmenting sporozoites within late oocysts (circled in the UMAP). **b**, Schematic representation of primary human hepatocyte infections with *Pf*SIP2 knockdown parasites. **c-d**, Knockdown by ATc (500 nM) withdrawal impaired (**c**) sporozoite invasion of primary hepatocytes at 3h pi (p=0.0147 and p=0.0009) and reduced (**d**) exoerythrocytic forms (EEF) formation at 2d pi (p=0.0048, p=0.0004, and p<0.0001) (Kruskal-Wallis test, Dunn’s correction). Data are represented as median. *n* indicates the number of independent wells across 3 biological replicates. *p<0.05, **p<0.01, ***p<0.001, ****p<0.0001. Schematics in **b** are adapted from ref. 36 under a Creative Commons licence CC BY 4.0.

*Pf*SIP2 expression is not detected in salivary gland sporozoites^5,21,38^ (although we cannot rule out expression below detection limits), suggesting that hepatocyte infection is a function of its expression in oocysts. Regardless, our data reveal that the downstream targets of *Pf*SIP2 could provide new candidates for transmission blocking strategies.

### Parasites interact with progenitor cells

Sequencing of both parasites and mosquito cells gave us the opportunity to identify possible interactions between the two organisms. We first analyzed the mosquito datasets from all four time points and the two treatments (ds*GFP* and ds*EcR*). After quality control using read and gene counts, mitochondrial read percentages, and complexity score (Extended Data Fig. 6a, Supplementary Table 1), we successfully profiled 55,789 high-quality midgut cells. Cells from each sample were then integrated using RPCA to correct for treatment and batch effects, followed by dimensionality reduction and clustering based on transcriptional profiles. We annotated the 16 resulting clusters based on marker gene expression and GO terms (Fig. 4a, Extended Data Fig. 6b, c, Supplementary Table 5, with a detailed description provided in the Supplementary Note). The major clusters included progenitor cells (intestinal stem cells/enteroblast, ISCs/EBs, headcase-positive); posterior enterocytes (pECs, nubbin-positive); anterior enterocytes (aECs, sugar transporter-positive); enteroendocrine cells (EEs, prospero-positive); visceral muscles (VMs, myosin-positive); and Proventriculus cells (also called cardia, PV, eupolytin-positive). A group of prospero-positive cells with mixed markers were annotated as EE-like. Interestingly, comparing ds*GFP* to ds*EcR* samples showed limited impact on mosquito transcriptomes, with fewer than 10 genes differentially expressed at each time point (Extended Data Fig. 6d, Supplementary Table 6). On the contrary, time post blood meal strongly affected gene expression across all major cell types (Extended Data Fig. 6e, Supplementary Table 7).

**Fig. 4.**
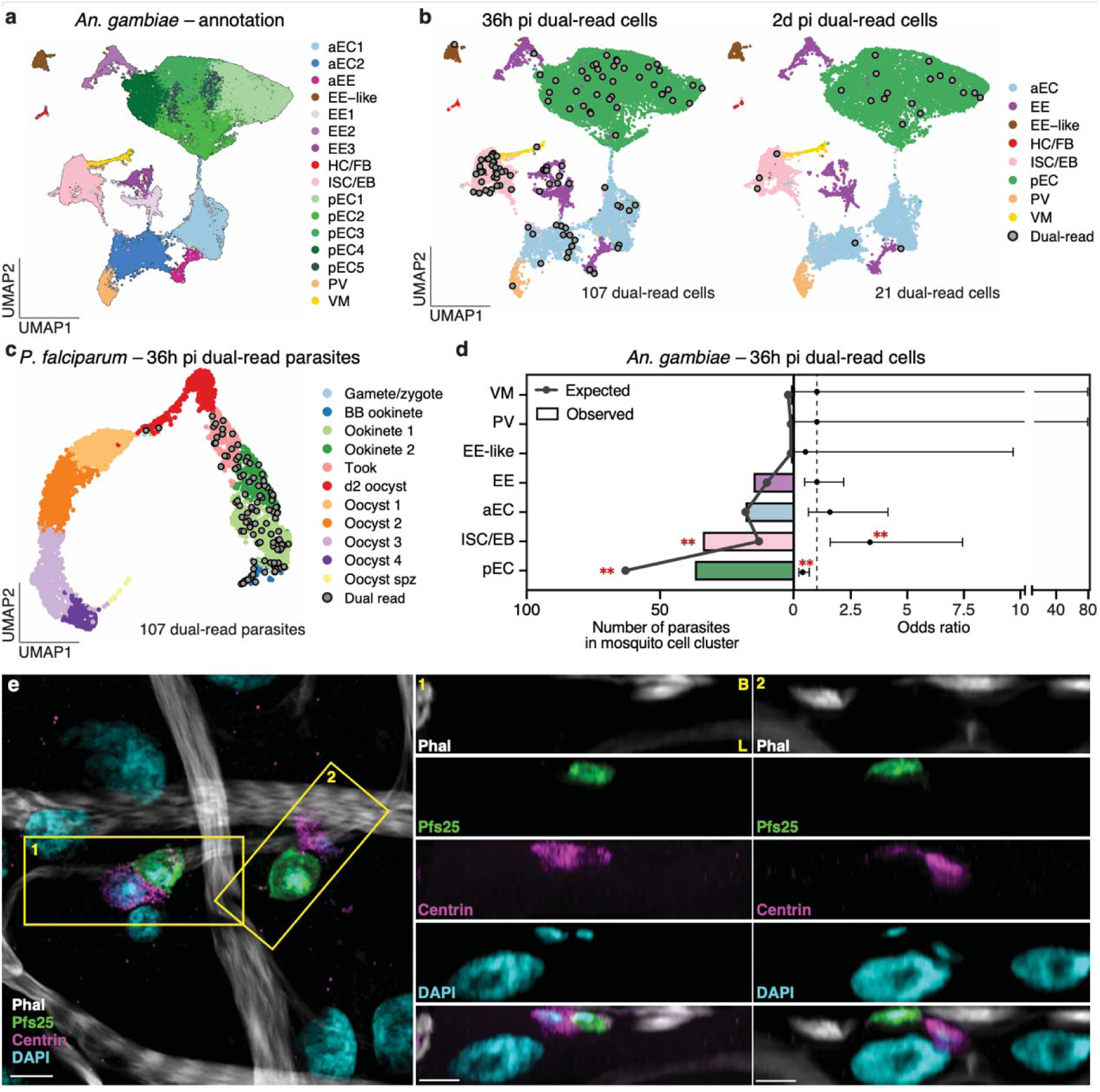
Parasite interactions with progenitor cells during midgut traversal. **a,** UMAP of *An. gambiae* midgut cells across all 4 time points forms 16 cell clusters, representing different cell types: anterior enterocytes (aECs), enteroendocrine cells (EEs), hemocyte and fat body cells (HC/FB), intestinal stem cells/enteroblasts (ISCs/EBs), posterior enterocytes (pECs), proventriculus (PV), visceral muscles (VMs), and a cluster with mixed markers annotated EE-like. **b,** UMAP plots of *An. gambiae* colored by cell type and overlaid with dual-read cells (cells with the same mosquito and parasite barcodes, indicated by grey circles) at 36h pi (left) and 2d pi (right). **c,** UMAP plot of *P. falciparum* from this study overlaid with dual-read parasites (grey circles) at 36h pi. **d,** Distribution of 107 dual-read parasites over eight independent samples in mosquito cell types at 36h pi with (left x axis) the observed vs. expected numbers under the random distribution and (right x axis) resulting odds ratio (point) ± 95% confidence interval (error bar; two-sided Fisher’s exact test with Benjamini-Hochberg (BH) multiple comparison correction, **: false discovery rate (FDR=0.003). **e**, Confocal imaging shows some parasites are inside of, or interact with, progenitor cells at 36h pi. Maximum intensity projection of a 2.31 µm Z-stack (0.21 µm between each plane) showing interactions between parasites (Pfs25, green) and progenitor cells (Centrin, magenta) (left image). Two orthogonal slices (yellow boxes) show parasites possibly inside (box 1) and adjacent (box 2) to progenitor cells. These interactions were observed across three independent infections. Nuclei were stained with DAPI (cyan), and muscle actin with phalloidin (Phal, white). LSM Plus processing was used. Scale bar: 5 µm.

To identify mosquito-parasite interactions, we focused on the 36h pi time point when parasites are crossing the epithelial layer. As ds*GFP* and ds*EcR* samples presented very similar mosquito and parasite transcriptomes, we pooled cells from these two groups (Extended Data Fig. 2a, 6d, Supplementary Table 6). We reasoned that a parasite captured inside or in strong association with a midgut cell would have the same cellular barcode as the mosquito cell. Despite the fleeting nature of crossing^11–13^, we detected 107 parasites sharing a barcode with a mosquito cell (defined here as dual-read cells) out of the 1,017 sequenced at this time point (10.5%) (Fig. 4b, Extended Data Fig. 7a). This percentage is significantly higher than the 3.9% observed at 2d pi (21 dual-read parasites out of 533 total, Fisher’s exact test, p-value <0.001), when parasites have mostly transformed into extracellular oocysts.

Interestingly, dual-read parasites appeared to interact preferentially with the ISC/EB midgut cluster (Fig. 4b-d). Indeed, 32.7% of dual-read cells (3.3% of total parasites) included a progenitor cell, although this cluster represents only 12.1% of all mosquito cells at this time point (Fig. 4d, OR 3.3, FDR = 0.003). This enrichment was not due to lower mRNA content in ISC/EBs, which might artificially increase parasite mRNA detection rates, and was not observed at 2d pi (Extended Data Fig. 7b, c). Parasites were also less likely to be associated with pECs than expected by chance (Fig. 4d, OR 0.37, FDR = 0.003). In differential expression analyses, no protein coding genes of midgut progenitor cells were affected by the presence of an interacting parasite (FDR = 1), and similarly, parasites interacting with a progenitor cell showed no difference in their transcriptomes relative to those of non-interacting parasites (FDR = 1), although these results may be due to the limited number of dual-read cells.

To confirm these findings, we performed confocal microscopy using an anti-centrin antibody that labels progenitor cells, as determined by their basal location, smaller triangular shape, and colocalization with the known progenitor marker armadillo^39^ (Extended Data Fig. 7d). We observed close interactions between parasites and progenitor cells, with 10.1% of parasites (35 out of 347) appearing to be inside or in close contact with an ISC/EB (Fig. 4e, box 1-2), or oriented towards one (Extended Data Fig. 7e, f). This number exceeds the 3.3% observed by scRNA-seq, suggesting that the single cell isolation process preferentially preserved the strongest interactions. Consistent with previous studies^12,15,40^, we also observed parasites near pECs that were either extruded or positive for the parasite surface protein Pfs25 (Extended Data Fig. 7e, f, Supplementary Video 1), potentially accounting for the reduced number of dual-read pECs associated with parasites.

We confirmed these findings by confocal microscopy in different mosquito-parasite combinations, observing close interactions between parasites and progenitor cells in 10.9% (56 out of 512) of parasites of the field isolate *P. falciparum* P5 in *An. gambiae* and in 8.71% (31 out of 356) of a different NF54 clone (referred to here as NF54_H) in a recently colonized *An. coluzzii* strain^41^ (Extended Data Fig. 7g, h).

Combined, these data reveal a preferential interaction, conserved across *Anopheles* species and *P. falciparum* isolates, between parasites and mosquito progenitor cells during traversal of the midgut epithelium, as ookinetes transform into oocysts.

### Late oocysts are associated with muscles

Interestingly, midgut progenitor cells were not observed to interact with ookinetes in the rodent-infecting *P. berghei* model but were instead associated with late oocysts, where their expansion eventually led to parasite death^42^. We, however, did not find evidence of accumulation of centrin-positive cells near oocysts at 7d pi (Fig. 5a), and on the contrary detected a decrease in the proportion of progenitors over time (Supplementary Table 5).

**Fig. 5.**
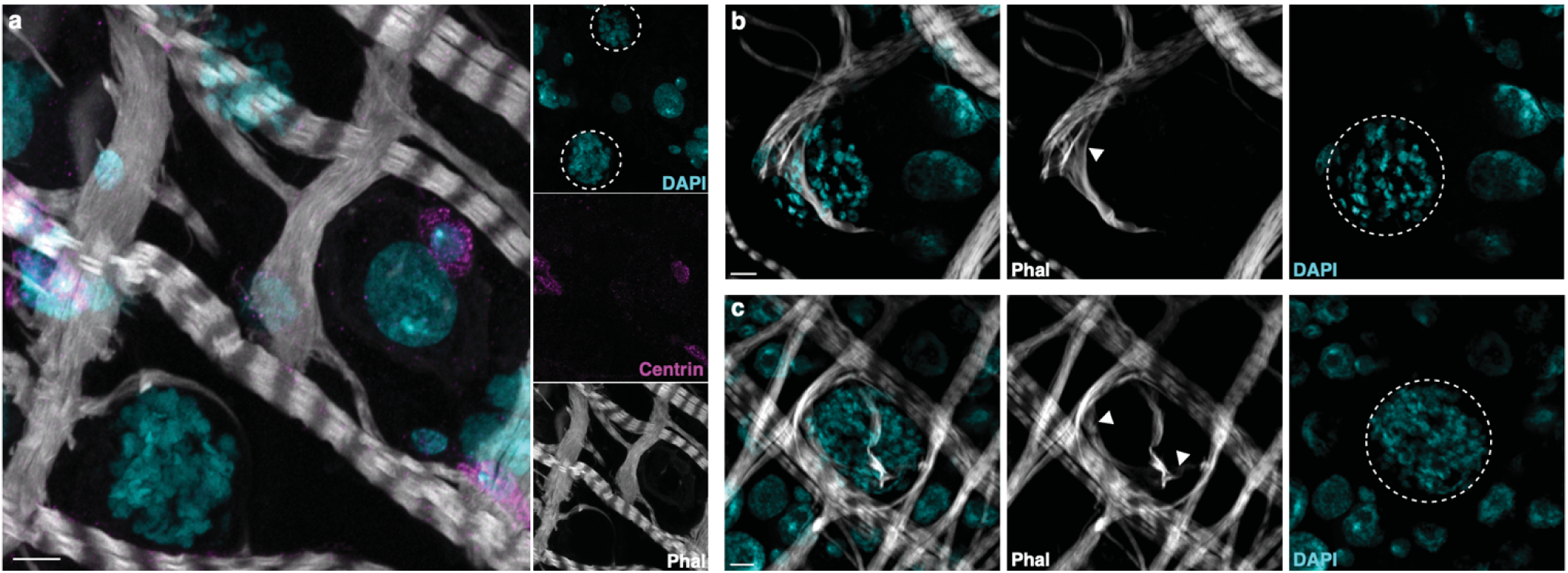
Late oocysts interact with visceral muscles. **a**, Maximum intensity projection of confocal images showing the absence of progenitor cell accumulation (centrin, magenta) around 7d oocysts (dotted circle in DAPI, cyan; Supplementary Video 2). **b-c,** two representative images showing muscle fibers (phalloidin, grey) (**b**) forcefully stretched (arrowhead) around a large oocyst (dotted white line) or (**c**) encasing an oocyst (Supplementary Video 3). Infections were performed in triplicate. Projections of 6.72 µm (a) and 2.94 µm Z-stack (b-c) are presented. Scale bar: 5 µm.

In our confocal microscopy analyses, however, we identified phalloidin-stained visceral muscles closely wrapped around late oocysts, with muscle fibers extending beyond their usual width (Fig. 5b, c; Supplementary Video 2-3). This interaction was conserved in several other mosquito-parasite combinations: *An. gambiae–P. falciparum* P5; *An. coluzzii*–NF54_H; and *An. stephensi*–ART29 (Extended Data Fig. 8a-c). Physical interactions with muscle cells may potentially tether oocysts to the midgut while minimizing disruption of the epithelial layer, as discussed later.

## Discussion

By focusing on developmental stages that had previously escaped large-scale analysis, our study provides a comprehensive reconstruction of the events characterizing the *P. falciparum* transmission cycle. This rich volume of information expands the repertoire of potential targets for preventing parasite transmission in strategies based on transmission-blocking drugs and vaccines^43^ or mosquito-targeted antimalarials^35,36^. Indeed, our functional analyses validate parasite factors (*Pf*ATP4, *Pf*LRS, and *Pf*SIP2) as essential for parasite growth and the acquisition of infectivity to human hepatocytes. We also reveal pathways potentially involved in the ookinete-oocyst transition, a possible role of *Pf*EMP1 in adherence of newly formed oocysts to the midgut, and processes fueling oocyst development, although these findings will require validation in future studies. Excitingly, our results demonstrate that much of the core machinery necessary for daughter cell formation (IMC, basal complex and other components) is conserved between blood stage parasites and oocysts, despite the sheer difference in scale of this process between these stages (tens of merozoites versus thousands of sporozoites).

The parallel sequencing of both parasites and mosquito cells allowed the identification of a previously unappreciated interaction between parasites and midgut progenitor cells. Ookinete traversal of the midgut epithelium is an asynchronous process that occurs between 24-36h pi. The best supported model for midgut crossing to date suggests that ookinetes initially invade an epithelial cell, then move laterally to adjacent cells before reaching the basal side, often leading to the extrusion of the invaded cells^14,44^. The observation that dual-read parasites contain not only ookinetes but also transforming tooks suggests that this interaction with progenitor cells occurs as parasites reach the basal side, where they transition into oocysts. Although our findings only begin to uncover the mechanisms governing ookinete exit and transformation into an oocyst, it is plausible that *P. falciparum* uses progenitor cells as landmarks for the basal side of the midgut. Here, parasite exit may be facilitated by the remodeling of tight junctions during division of ISCs^45,46^, although several alternative hypotheses are possible. It is likely that these interactions occur after ookinetes have rapidly escaped extruding pECs, as observed both in this study and in several others^14,15,40^ and in alignment with our observation of a slightly reduced number of parasite-associated pECs.

In our analysis, *P. falciparum* oocysts were found to interact tightly with muscle cells. This interaction may be the result of the specific coevolutionary trajectories of human malaria parasites with human-biting mosquitoes. Indeed, *P. falciparum* possesses mechanisms to minimize the damage inflicted to its *Anopheles* vectors^24,47^, including a low number of ookinetes crossing the midgut epithelium. Rodent malaria parasites, on the other hand, have high parasite loads that induce substantial epithelial damage and overall fitness costs^48,49^, and their oocysts are deeply embedded in the epithelium^50^ likely causing damage and consequent ISCs proliferation^42^. A close association with muscle cells might allow *P. falciparum* oocysts, which on the contrary jut farther out of the midgut^50^, to limit epithelial damage by tethering growing parasites in a less invasive way. Further research will clarify the biological relevance of the mosquito-parasite interactions we unveiled in this study.

## Methods

### Rearing of *Anopheles* mosquitoes

*Anopheles gambiae* (G3 strain), *An. stephensi* (SDA-500), and recently colonized *An. coluzzi*^41^ mosquitoes were reared in an insectary maintained at 27°C, 70-80% relative humidity, on a 12h:12h light:dark cycle. Larvae were fed on fish food and adults were provided with 10% w/v glucose solution *ad libitum*. Adult mosquitoes were fed with purchased human blood (Research Blood Components, Boston, MA) to induce egg laying.

### dsRNA production and microinjection

PCR fragments of *eGFP* control (495 bp) and *EcR* (AGAP029539; 435 bp) were amplified from the plasmids pCR2.1-eGFP and pCR2.1-EcR as previously described^52^. PCR product was used to transcribe dsRNA using the Megascript T7 transcription kit (Thermo Fisher Scientific), per manufacturer’s protocol. The resulting dsRNA was purified via phenol-chloroform and diluted to a concentration of 10 µg/µL. Microinjections were performed as detailed previously^24^, with adult mosquitoes being injected one day post-eclosion and given an infectious blood meal 3 days post-injection.

### EcR knockdown confirmation

Resulting knockdown confirmation post-injection was performed as previously described^24^. Pools of 6-8 infected female mosquitoes were aspirated at 24h pi and transferred to TRI reagent (Thermo Fisher Scientific). RNA was extracted, DNase treated, quantified, and reverse transcribed. cDNA from four biological replicates was diluted tenfold and run in duplicate for qRT-PCR. Gene-specific primers were used, and relative quantification was performed with the ribosomal gene *Rpl19* as the reference (Supplementary Table 8)^53^. RNA extraction from one replicate of the single-cell experiments failed and were thus redone with samples collected at 2d pi. For phenotypic EcR knockdown confirmation, individual mosquito ovaries were also collected 7d pi in 80% ethanol for later manual egg counts.

### Culturing of *P. falciparum* parasites

*P. falciparum* strains used in this study include: (1) NF54 originally provided by C. Barillas-Mury from BEI Resources (MRA-1000), (2) NF54_H, highly infectious to mosquitoes, from Michael Delves’s laboratory, (3) an artemisinin-resistant, Cambodian field isolate ART29^54^, and (4) a Burkina Faso polyclonal isolate P5^25,35^. Asexual stages of parasites were cultured at 37°C between 0.2 and 2% parasitemia in human erythrocytes at 5% hematocrit (Interstate Blood Bank, Memphis TN) using RPMI medium 1640 supplemented with 25mM HEPES, 10mg/l hypoxanthine, 0.2% sodium bicarbonate, and 10% heat-inactivated human serum (Interstate Blood Bank) under a gas mixture of 5% O_2_, 5% CO_2_, balanced N_2_. Stage V female and male gametocytes were induced by raising parasitemia over 4% and incubating cultures for ~14–20 days with daily media change.

### *P. falciparum* infections of *Anopheles* mosquitoes

Female mosquitoes were blood fed on a mix of *P. falciparum* gametocyte culture, human red blood cells and human serum (ratio 1:3:6) via heated membrane feeders 3-5 days post eclosion, and introduced into a custom-built glove box (Inert Technology, Amesbury MA) as previously described^24^. Unfed mosquitoes were removed, and remaining mosquitoes were kept on 10% glucose solution *ad libitum*, with added ATc or compound (see below). At dissection time points, mosquitoes were aspirated into 80% ethanol, frozen for 10 minutes, and transferred to phosphate-buffered saline (PBS) on ice.

### Generation of single-cell suspension

Forty midguts from *P. falciparum*-infected mosquitoes injected with ds*GFP* or ds*EcR* were dissected at 36h, 2d, 4d, and 7d pi across four biological replicates in ice-cold PBS. Any remaining blood meal was removed before storing the dissected midguts in ice-cold PBS. Dissections were completed within 1 hour to preserve the viability of parasite and mosquito cells. Midguts were pooled into 200 µl of PBS with 1 mg/mL elastase (Sigma-Aldrich E7885) and 1 mg/mL collagenase IV (Thermo Fisher Scientific 17104019), and incubated at 30°C, 300 rpm for 30 min in a shaking heat block. To facilitate the isolation of single cells, samples were disrupted every 10 min by pipetting with low protein-binding tips.

Samples were assessed using trypan blue staining to quantify live, single midgut cells during protocol optimization. This included systematically testing variables such as digestion time, temperature, tissue preparation (intact or minced gut), and types of pipette tips. On average, approximately 400 live, single cells were consistently recovered per mosquito midgut, while many cells remained in clumps. Given the low number of parasites (~20-30) relative to mosquito cells (~5,000) in a midgut, rather than further optimizing for complete dissociation of the midgut, we chose to enrich the parasite-to-mosquito cell ratio by removing cell clumps as they were composed mostly of midgut cells. Therefore, the cell suspension was sequentially filtered through 70 µm, 40 µm, and 20 µm cell strainers (pluriSelect), with each strainer being immediately washed with 200 µl of 1x PBS to collect any adherent cells. The 20 µm cell strainers were not used for the 7d pi samples, as oocysts at this time point are, on average, over 25 µm in diameter^24^. Based on the final dataset, this approach yielded an approximate 1:16 parasite-to-mosquito cell ratio. Given an estimated 5,000 midgut cells and 30 oocysts per gut – an expected ratio of about 166:1 – our protocol represents a 10-fold enrichment of parasites.

We chose not to collect samples beyond 7d pi, as oocysts larger than 30 µm may clog the 10X Chromium chip. We chose not to use fluorescence-activated or magnetic-activated cell sorting as they both lengthen the isolation protocol and cause stress, reducing cell viability. The final single-cell suspension was pelleted, resuspended in 20 µl of PBS, and cell concentration and viability were determined by hemocytometer count with trypan blue staining.

### Single-cell library preparation and sequencing

Single-cell libraries were prepared following the Chromium Next GEM Single Cell 3’ Reagents Kit v3.1 (Dual Index) User Guide (RevC). Briefly, single-cell suspensions were diluted to target a recovery of 4,000 cells per sample, then mixed with reverse transcription reagents and barcoded gel beads. The mixtures were loaded into wells containing partitioning oil on a 10X Chromium chip to generate single-cell emulsions. Samples from ds*GFP* and ds*EcR* mosquitoes collected at the same time point were loaded into two separate wells on the same chip. In total, 16 chips were used across four time points and four biological replicates. Within each emulsion droplet, individual cells were lysed, mRNA molecules were captured, and cDNA were generated with cell-specific barcodes. Barcoded cDNA were pooled, then amplified, fragmented, and purified using SPRIselect, followed by ligation with Illumina adaptors and sample-specific barcodes. Each sample was assessed using an Agilent Bioanalyzer to ensure library integrity and quantified by qPCR using i7 and i5 Illumina primers, following the protocol from Universal Kapa Library Quantification Kit (Roche). The cDNA libraries from each sample were then mixed in equal proportions, spiked with 1% phage cDNA (PhiX, Illumina). The proportion of parasite reads relative to mosquito reads was estimated using an initial Illumina iSeq 100 run, which showed that around 0.05%, 0.1%, 0.6%, and 1.3% reads mapped to parasites at 36h, 2d, 4d and 7d pi, respectively. To obtain sufficient coverage of parasites transcripts, final sequencing was performed across all lanes of two NovaSeq S4 flow cells (Broad Institute), with an expectation of 1,974–51,316 reads per parasite ranging from 36h to 7d pi (see supplementary note for detailed calculation).

### scRNA-seq data processing and analysis of *P. falciparum*

FASTQ files from each sample originating from different flow cells were concatenated and mapped to the genomes of *P. falciparum* 3D7 (PlasmoDB.org, version 58)^55^ using the 10X Cell Ranger software v7.0.1^56^. The resulting count matrix for each sample was processed and filtered using Scanpy (v1.9.1) in Python (v3.10)^57^. Based on the individual sample profile, low-quality cells, dead cells, and empty droplets were removed according to the number of reads (UMI count), number of genes (gene count), and proportion of mitochondrial reads per cell (Supplementary Table 1). The count matrix from each sample was merged into a single AnnData object by Scanpy, and doublets were removed running Scrublet software in Python before conversion to Seurat using SeuratDisk’s function Convert^58^. After quality control, the average number of reads per cell ranged from 649 at 36h pi to 9,043 at 7d pi. Pseudobulk differential expression analyses were initially conducted treating the samples as independent observations for comparison. The pseudobulk count matrix was generated for each sample by summing raw gene counts across all cells in Python with Pegasus. Several comparisons were conducted – between each time point in parasites from ds*GFP*-injected mosquitoes as well as between parasites from ds*EcR*- and ds*GFP*-injected mosquitoes of the same time point – using a Python wrapper for the DESeq2 package from R^59^. Significance was set as log_2_ fold change greater than 1 or less than −1, and an adjusted p-value below 0.05. Functional enrichment analysis of upregulated or downregulated gene lists was performed using GoProfiler^60^.

A standard single cell analysis pipeline was then performed with Seurat and dependent packages in R 4.3.2 in RStudio 2023.9.1.494^61^. Highly variable genes were identified with the FindVariableFeatures function before data were scaled (ScaleData, default settings) and a Principal Component Analysis (PCA) plot was generated based on these features only. Next, high-quality parasite cells were integrated with existing datasets, including blood bolus ookinetes at 24h and 48h pi, gametes/zygotes/ookinetes from 2-20h pi, and sporozoites released from oocysts at 12d pi^21,22^. Briefly, Seurat objects (v5) were created for external datasets and reads were normalized to 10,000 transcripts, before the three Seurat objects were merged. Variable features were identified with default parameters of the FindVariableFeatures function, the data was scaled (ScaleData), and PCA was performed on the first 50 dimensions (RunPCA, default settings). Integration to correct for batch effect and differences in single cell methodology was conducted using reciprocal PCA (RPCA), which is the preferred approach when minimal overlap between datasets is expected. Subsequently, a UMAP was generated including the top 12 principal components through the RunUMAP function, with parameters set to min.dist=0.4, and repulsion.strength=2. Cell clusters were defined by the Louvain algorithm at a resolution of 0.5. Cluster markers were identified using the FindAllMarkers function, focusing only on positive markers with a log_2_ fold change above 1. Genes with false discovery rates below 0.01 were utilized to label each cluster.

The Seurat object was converted into a cds object in Monocle3^62^, and the trajectory analysis was performed using the learn_graph function (close_loop = F, ncenter=500). Pseudotime analysis was conducted by setting the node in the gamete/zygote cluster as the start point. Gene network analysis focused on our data and was performed using the graph_test function (neighbor_graph=”principal_graph”), to identify co-expressed genes that change as a function of pseudotime (FDR < 0.05). Co-expressed genes were further clustered by the find_gene_modules function with a resolution of 0.003. The expression of all genes within each gene cluster was aggregated using the aggregate_gene_expression function. Enriched gene ontology terms were determined by analyzing all the genes within a gene cluster in GoProfiler^60^. Differential expression of protein coding genes was compared between dual-read parasites at 36h pi and parasites that do not share a barcode with a mosquito cell, binning for parasite cluster, using logistic regression. A subsequent analysis focused on dual-read parasites only at 36h pi, and compared those interacting with an ISC/EB to the remainder.

### Mosquito scRNA-seq data processing and analysis

All FASTQ files from the different sequencing runs were concatenated and mapped to the *An. gambiae* PEST genome (VectorBase.org, version 58)^55^, using the 10X Cell Ranger software v7.0.1^56^. Quality control was initially performed in Python using Scanpy based on the number of reads (UMI count), number of genes (gene count), percentage of reads mapping to the mitochondrial genome and cell complexity (Supplementary Table 1). Cells with a complexity score (ratio of number of transcripts by genes logged) below 0.7 or more than 30% reads mapping to the mitochondrial genome were removed^63–68^. The mitochondrial threshold was chosen based on published scRNA-seq studies of *Anopheles* hemocytes^63^ and *Aedes* midguts^64^, noting that a similar threshold (25%) was used by two scRNA-seq studies on the midgut of *Drosophila melanogaster*^65^ and *Culex tarsalis*^66^, whereas snRNA studies used lower cutoffs due to inherent differences in the technology^67,68^. The resulting count matrix from each sample was merged into a single AnnData object by Scanpy, before conversion to Seurat using SeuratDisk’s function Convert. Doublet removal was performed in R using scDblFinder^69^. Quality metrics indicate that mitochondrial percentage is lower in 36h and 2d pi samples compared to the 4d and 7d ones, likely due to a combination of biological factors, including time after blood feeding, age, and infectious stages (ookinetes/young oocysts versus large oocysts).

After quality control, mosquito cells from each time point were processed using a similar workflow as for parasite analysis. The top 25% variable genes were identified, the data was scaled, and PCA was performed computing the first 100 PCs. Integration was conducted using RPCA with the highest k.weight (64) to correct batch effect while avoiding overcorrecting for biological variation between time points and treatments. The UMAP was calculated using the first 56 PCs, and cluster identification was based on the Louvain algorithm with a resolution of 0.3. Positive markers were identified using the FindAllMarkers function with a log_2_ fold change greater than 1. Proventriculus, anterior, and posterior midgut scores were calculated based on *An. gambiae* midgut bulk RNA-seq results^51^. Clusters were then annotated by identifying the functional enrichment of each cluster’s marker genes as well as cross-referencing cluster markers from four references: the Fly Cell Atlas, the *Drosophila melanogaster* gut single-cell study, the *Aedes aegypti* gut single-cell study, and the hemocyte single-cell study from *An. gambiae*^63,65,67,68^. *An. gambiae* orthologs were identified using VectorBase^55^, and all significant cluster markers with an ortholog specific to a cell type were used for cluster annotation.

Pseudobulk analyses were performed to assess the effect of either *EcR* knockdown or time post infectious blood meal on the midgut transcriptome using AggregateExpression from the Seurat package. To assess the effect of *EcR* knockdown on transcription, differential expression analyses were performed using DESeq2^59^ by pooling cells from each cell type at each time point. To evaluate the effect of time on transcription, only cells from ds*GFP* mosquitoes were used to compare between consecutive time points. Functional enrichment analyses were performed using GoProfiler^60^ when more than 20 genes were significant. Dual-read differential expression analyses of protein coding genes were performed using logistic regression framework comparing dual-read cells at 36h pi with cells not interacting with a parasite at the same timepoint, binning for dsRNA treatment and annotation. The analysis was repeated by comparing protein coding gene expression between dual-read ISC/EB cells at 36h pi and ISC/EBs not interacting with a parasite.

### Post-infection exposure assays via sugar feeding

Candidate target selection for drug exposure was based on two criteria: (1) belonging to a distinct gene cluster in our gene model, and (2) having known potent activity against the candidate target in the *P. falciparum* blood stage. Four drugs were selected: cipargamin (ChemPartner), MMV670325, MMV643121 and MMV007839 (Medicine for Malaria Venture)^70–73^. Stock solutions were made in DMSO at 100 mM, stored at −20°C and diluted 1:1,000 in 10% glucose solution shortly before sugar change. As the bioavailability of these compounds in mosquitoes is unknown, a single high concentration of 100 µM was chosen initially as it is close to the solubility limit while keeping DMSO content to 0.1%. Mosquito compound exposure through sugar feeding was performed similarly as previously described^54^. Briefly, sugar solutions containing compounds or corresponding controls (0.1% DMSO) were replaced every day from 2d to 10d pi (unless specified). For oocyst counts, midguts were collected at 7d pi and stained in 0.2% (w/v) mercurochrome (Sigma-Aldrich, St. Louis, MO). Midguts were imaged using an Olympus Inverted CKX41 microscope, and oocyst number and mean size were calculated using OocystMeter^74^. For salivary gland sporozoite quantification, salivary glands of individual females were dissected at 14d pi in PBS and counted on a hemocytometer.

### Plasmid construction

The *Pf*SIP2 homology-directed repair (HDR) plasmid was assembled by Golden Gate cloning (New England Biolabs) using purified PCR products of (1) a C-terminal homology region (Cterm-HR) that was codon-altered to avoid repeated CRISPR cuts, (2) a cassette containing hemagglutinin (HA)-tags, tet-aptamers^75^, a TetR-DOZI fusion protein^75,76^, and a human dihydrofolate reductase (hDHFR) selection marker, (3) a 3’ HR, and (4) a bacterial backbone. Primers used to amplify fragments are listed in Supplementary Table 8 and assembly was confirmed by whole plasmid sequencing (Plasmidsaurus). Two CRISPR-Cas9 guide plasmids were constructed to cut the C-terminus of the endogenous *Pf*SIP2. Guide oligos were annealed and ligated into BpiI-digested pRR216^77^, which contains *Sp*Cas9, a U6 guide cassette, and a yeast dihydroorotate dehydrogenase (DHODH) selection marker. Sequences were confirmed by Sanger sequencing (Psomagen).

### Generation of *Pf*SIP2-cKD parasite line

The HDR and guide plasmids of *Pf*SIP2 were extracted using a Qiagen Maxi kit, and 100 μg of the HDR plasmid was linearized with XhoI, NotI, and ApaL1 at 37°C overnight. The linearized HDR plasmid and a mixture of the 2 guide plasmids (60 μg each) were sterilized by ethanol precipitation and resuspend in TE buffer. The plasmids were then mixed with 270 µl of Cytomix (120 mM KCl, 0.15 mM CaCl_2_, 2 mM EGTA, 5 mM MgCl_2_, 10 mM K_2_HPO_4_, 10 mM KH_2_PO_4_, 25 mM HEPES, pH 7.6) and transfection by red blood cell loading was performed as described^78^. The parasite line was maintained at 5% hematocrit with 500 nM ATc (Sigma) from the onset of transfection. Drug pressure for the guide plasmid was applied from 6 hours to 5 days post-transfection using 1.5 μM DSM1 (Sigma), whereas drug pressure for the HDR plasmid was maintained with 2.5 nM WR99210 hydrochloride (Sigma) starting 6 hours post-transfection until integration was confirmed. To do so, genomic DNA was extracted from WT and PfSIP2-cKD lines using the Qiagen Blood & Tissue kit. Each integration site of the double crossover was amplified as well as the whole locus and PCR products were analyzed by gel electrophoresis.

### Asexual stage growth assay

ATc was washed out from PfSIP2-cKD parasites three times with RPMI1640. The resulting parasites were diluted to 0.5% starting parasitemia at 2.5% hematocrit and treated with either 500 nM ATc, 1.5 nM ATc, or 0.025% of DMSO. Parasitemia was calculated daily by counting Giemsa-stained smears of each technical replicate. The experiment was repeated twice, with 4 technical replicates for each biological replicate. Data were analyzed by two-way ANOVA with Dunn’s correction.

### Mosquito infection with transgenic parasites

Transgenic parasites were cultured in the asexual blood and gametocyte stages and fed to mosquitoes as described above, with the following specifications for ATc usage. Both asexual and gametocyte cultures were maintained in 500 nM ATc (dissolved in DMSO). One hour prior to mosquito infection, ATc was washed out from the gametocyte cultures by replacing the media three times, and either 500 nM ATc or 0.025% DMSO was added to the blood meal. Mosquitoes were maintained on a 10% glucose solution supplemented with 100 µM ATc, which was dissolved directly in the solution.

### Culturing of primary human hepatocytes and infection with transgenic parasites

Cryopreserved human primary hepatocytes (BioIVT, Westbury, NY) were thawed, seeded on collagen-coated micropatterned islands in 96 well plates, and infected with WT or *Pf*SIP2-cKD sporozoites as previously described^79^. In short, WT or *Pf*SIP2-cKD-infected mosquito salivary glands were dissected in Schneider’s medium (Gibco) containing 200 U/mL of penicillin, 200 µg/mL of streptomycin (2X, Gibco) for less than 1 hour. Sporozoites were released from glands, filtered through a 35 µm cell strainer and counted on a hemocytometer. Parasites were spun at 10,000 *g* for 3 min, resuspended in hepatocyte media containing 2.5 μg/mL Fungizone (Cytiva) and 2X penicillin/streptomycin as well as either 500 nM ATc or 0.025% of DMSO and seeded onto hepatocytes. For invasion assays, 7,000 sporozoites were seeded per well, wells were fixed in 4% paraformaldehyde (PFA) 3h pi and in and out *Pf*CSP staining was performed^79^. For infection assays, 70,000-100,000 sporozoites were used, and subsequently washed off 3h pi before supporting cells from the male mouse fibroblast cell line 3T3-J2 (Rheinwald and Green, 1975) were added. Cells were washed daily, and wells were fixed at 2d pi in 100% methanol and stained with rabbit αPfHSP70 monoclonal primary antibody (antibodies-online, ABIN361730) and goat anti-rabbit IgG 546.

### Immunofluorescent microscopy

Midguts were dissected from females at specified time points. Any remaining blood bolus was removed in ice-cold PBS when present and clean midguts were incubated at room temperature in 4% PFA for 45 min, followed by 3 washes in PBS for 10 min each. Midguts were permeabilized and blocked in 0.1% Triton X-100, 3% bovine serum albumin (BSA) in PBS for 1 hour at room temperature or overnight at 4°C before incubation in primary antibody at room temperature for 2 hours, shaking. The following primary antibodies were used: 1:200 anti-*Toxoplasma gondii* centrin-1 rabbit polyclonal (Kerafast EBC004); 1:5 anti-*Drosophila* armadillo mouse monoclonal supernatant N2 7A1^80^ (deposited to the DSHB by E. Weischaus), 1:200 anti-Pfs25 mouse monoclonal 4B7^81^ (deposited to BEI resources by Louis H. Miller and Allan Saul). Another 3 washes in PBS were performed before incubating the samples in secondary antibodies and phalloidin for 1 hour at room temperature rocking (goat anti-rabbit IgG AlexaFluor 488, goat anti-mouse IgG AlexaFluor 488, goat anti-mouse AlexaFluor 568 from Invitrogen used at 1:400, Ebioscience Phalloidin eFluor 660 used at 1:300). Midguts were then stained in PBS containing 5 µg/mL DAPI before the remaining 3 washes in PBS only. Midguts were mounted using VECTASHIELD HardSet Antifade Mounting Medium and imaged on a Zeiss Inverted Observer Z1 with Apotome3 or a Zeiss LSM 980 confocal microscope. Using ZEN 3.10 software, confocal images were processed with LSMPlus while Airyscan images were processed with default parameters, followed by brightness and contrast adjustments in Fiji (version 2.14.0). Gamma was not adjusted unless specified.

### Statistical analysis

Specific statistical tests are detailed in the corresponding figure legend. Confirmation of EcR knockdown and analysis of parasite prevalence, intensity and infection rates were conducted using GraphPad Prism 10.1.1. Odds ratios of dual-read parasites in mosquito cell types were calculated in R using Fisher’s exact tests followed by correction for multiple comparisons with BH.

## Acknowledgements

The authors thank Rhoel Dinglasan and Marc-Jan Gubbels for insights on the centrin antibody, Jeffrey D. Dvorin for the tetR-DOZI and guide plasmids, and Michael Delves for the NF54_H parasites. We are grateful to Scott E. Lindner for valuable discussions on rRNA, to Kirk W. Deitsch for insights into PfEMP1, and to the members of the Catteruccia laboratory for comments. We thank Lucia Ricci for help with graphics, and we acknowledge Neha Damaraju for her assistance with hepatocyte infections. The armadillo monoclonal antibody N2 7A1 developed by E. Wieschaus was obtained from the Developmental Studies Hybridoma Bank, created by the NICHD and maintained at The University of Iowa. We are deeply grateful to VEuPathDB for aiding the analysis of the datasets. FC is funded by the Howard Hughes Medical Institute (HHMI) as an Investigator and by the National Institute of Health (NIH) grants R01AI148646 and R01AI153404. LEdV was funded by a Rubicon grant (452021309) from the Dutch Research Council (NWO).

## Author Contribution statement

YY, EC, WRS, DP, and FC conceived the study. YY, EC, ED, and ASP performed the dissections for the scRNA experiments, and YY and EC performed the single cell isolation and scRNA library generation. YY, LHV, and DP performed the scRNA analysis. YY, LHV, EC, FA, and TAR performed and analyzed drug infection experiments. YY, ED, and LEdV generated transgenic parasites. YY performed *Pf*SIP2-cKD experiments and YY, LHV, SM, and JK performed the primary human hepatocytes infections. EN reared mosquitoes used in the study. NS generated gametocyte culture and infected mosquitoes. LHV, EC, and ARC performed immunofluorescence imaging and analysis. SNB, DP, WRS, and FC provided supervision and oversaw experiments and analyses. YY, LHV, EC, and FC wrote the original draft. YY, LHV, WRS, and FC reviewed and edited the manuscript. All authors approved the final manuscript.

---

**Extended Data Figure 1.**
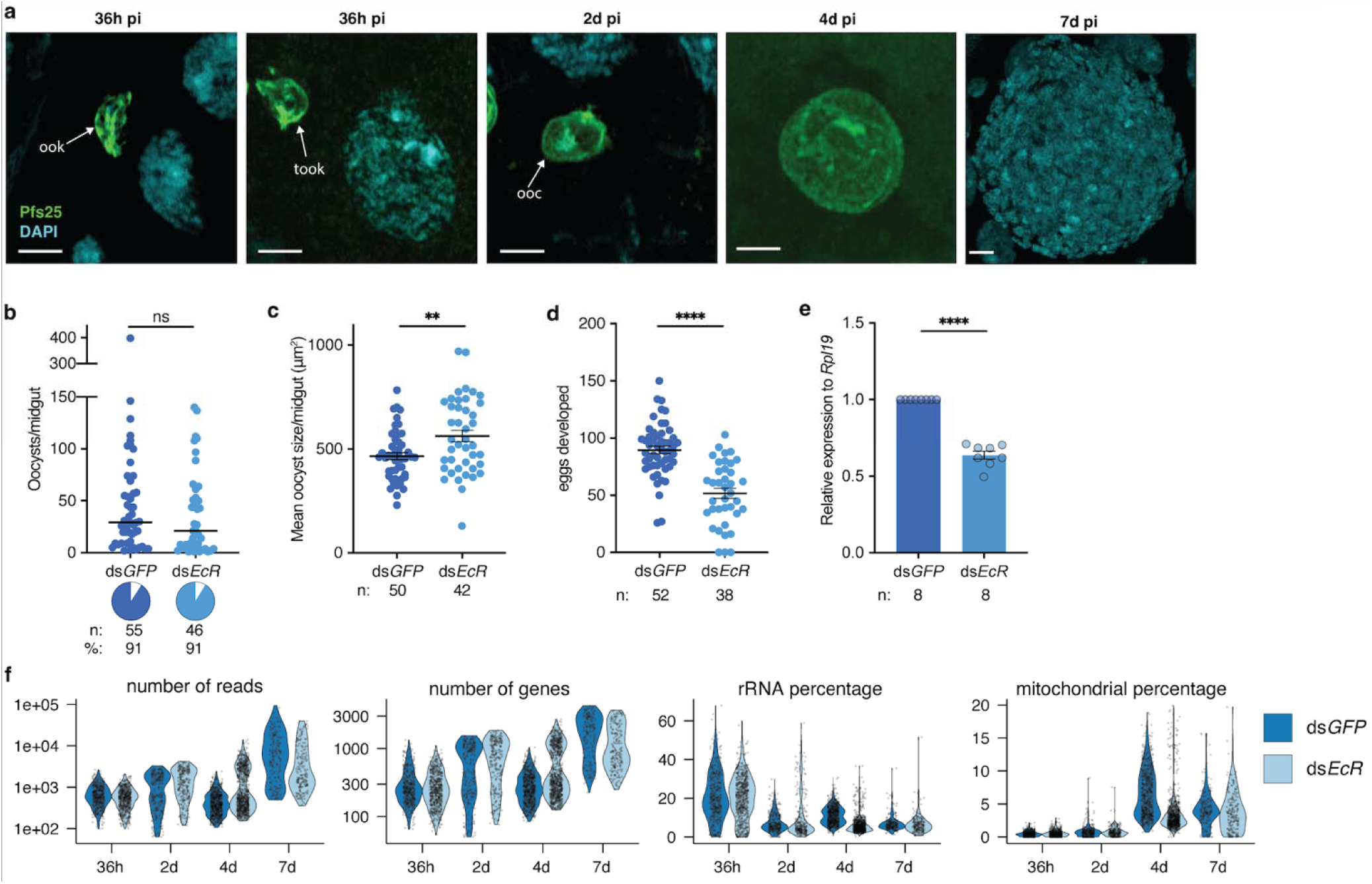
Quality control (QC) of scRNA-seq in two metabolic conditions. **a**, Representative images of the parasite forms collected during the 4 experimental time points. Ookinetes (ook) and tooks found at 36h pi, and oocysts (ooc) identified at 2d, 4d, and 7d pi. Parasites were stained with Pfs25 (green) and nuclei with DAPI (cyan). Scale bar: 5 µm. **b-c**, *EcR* knockdown across 4 replicates (**b**) did not affect oocyst prevalence (pie-charts, Fisher’s exact test, two-tailed) or intensity (Mann Whitney test, two-tailed), but (**c**) increased oocyst size at 7d pi (Mann Whitney test, two-tailed). **d**-**e**, Knockdown of *EcR* was confirmed by (**d**) detecting the expected reduction in egg numbers in the ovaries (unpaired t-test, two-tailed) and (**e**) RT-qPCR (unpaired t-test, two-tailed). **f**, Violin plots displaying key metrics of the *P. falciparum* scRNA-seq data after QC. *n* indicates the number of mosquitoes, % the prevalence of infected mosquitoes. Data are represented as mean ± SEM in panels **c**, **d** and **e**, and as median in panels **b**.

**Extended Data Figure 2.**
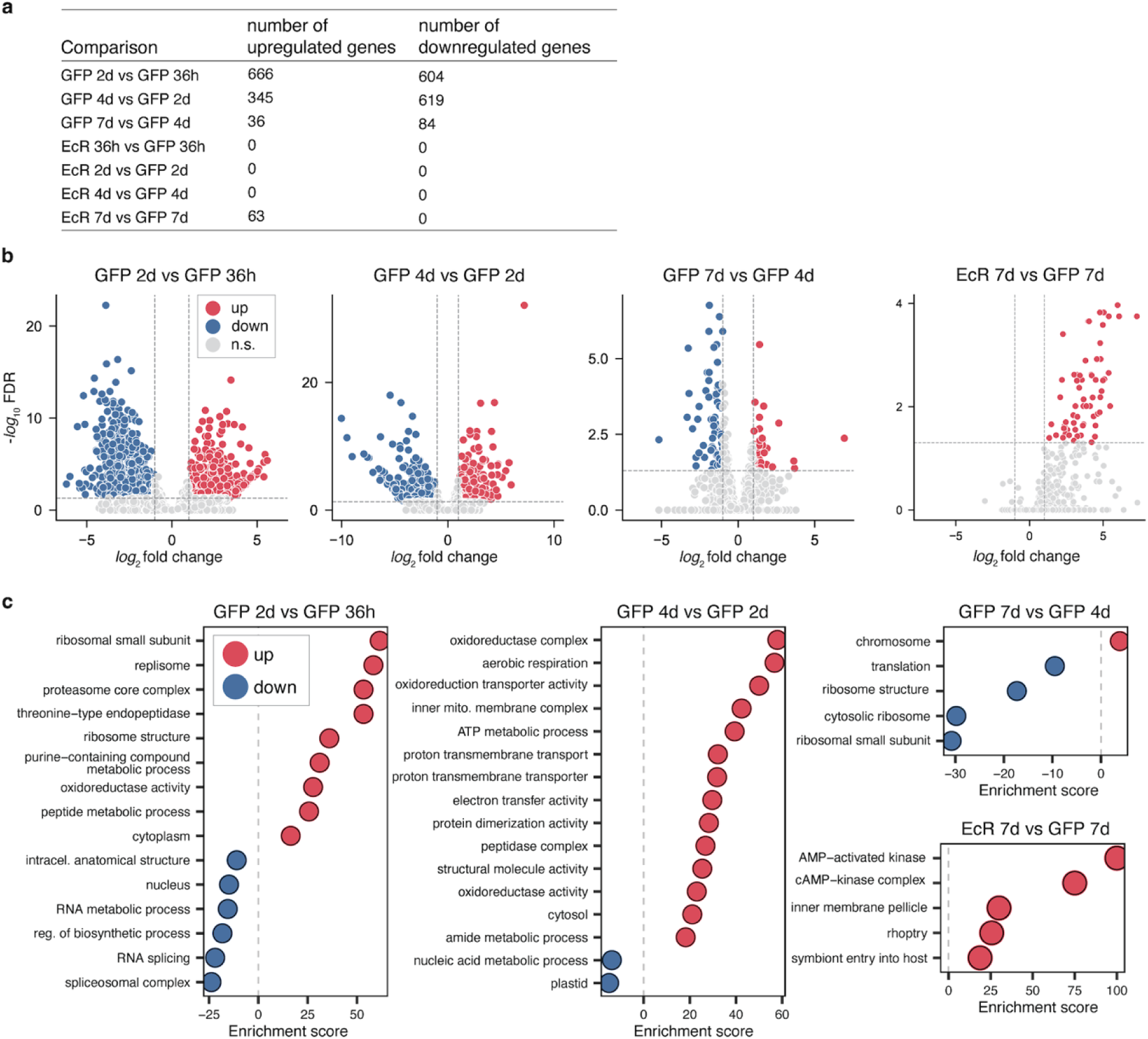
Pseudobulk differential expression analysis of *P. falciparum* across time and treatment. **a**, Table summarizing the number of up- and down-regulated genes identified in the comparisons of time points and/or treatments by pseudobulk (significance defined as FDR<0.05 and absolute value of log_2_ fold change>1). **b**, Volcano plots representing the differentially regulated genes in the four comparisons that yield significant changes. **c**, Corresponding GO terms that are significantly enriched in up- and down-regulated genes (Fisher’s one-tailed test, g:SCS algorithm for multiple test correction).

**Extended Data Figure 3.**
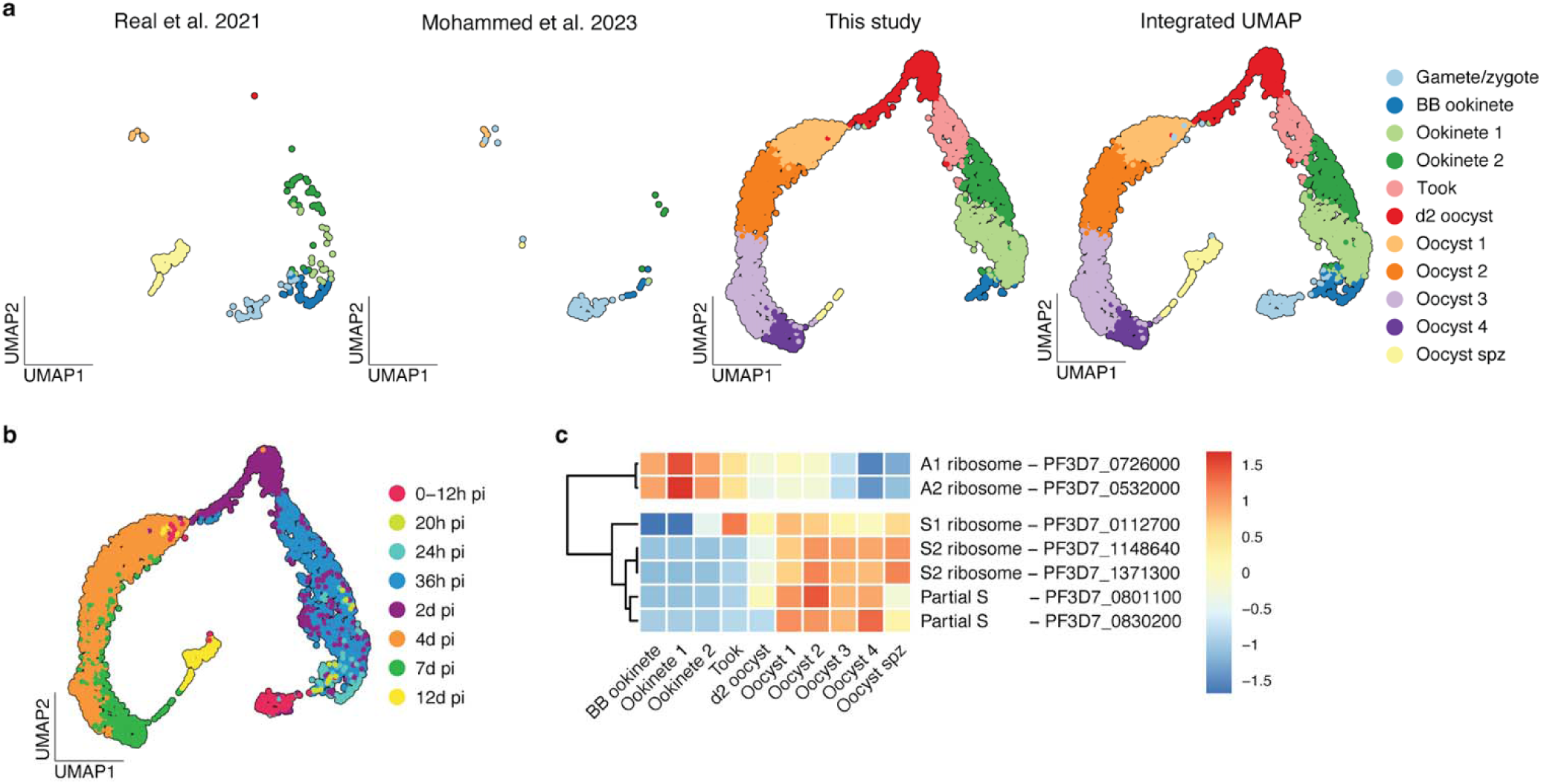
Integrated *P. falciparum* UMAPs across data origin and time. **a-b**, UMAP plots representing the integrated data of *P. falciparum* (**a**) split by their study of origin and (**b**) colored by time points of single cell collection. **c,** Heatmap showing the Z scores of expression patterns of ribosomal RNA (rRNA). Each row represents a cumulative expression of 18S (when present), 5.8S and 28S for each rRNA type (A1, A2, S1, S2). The gene ID given is of the corresponding 28S.

**Extended Data Figure 4.**
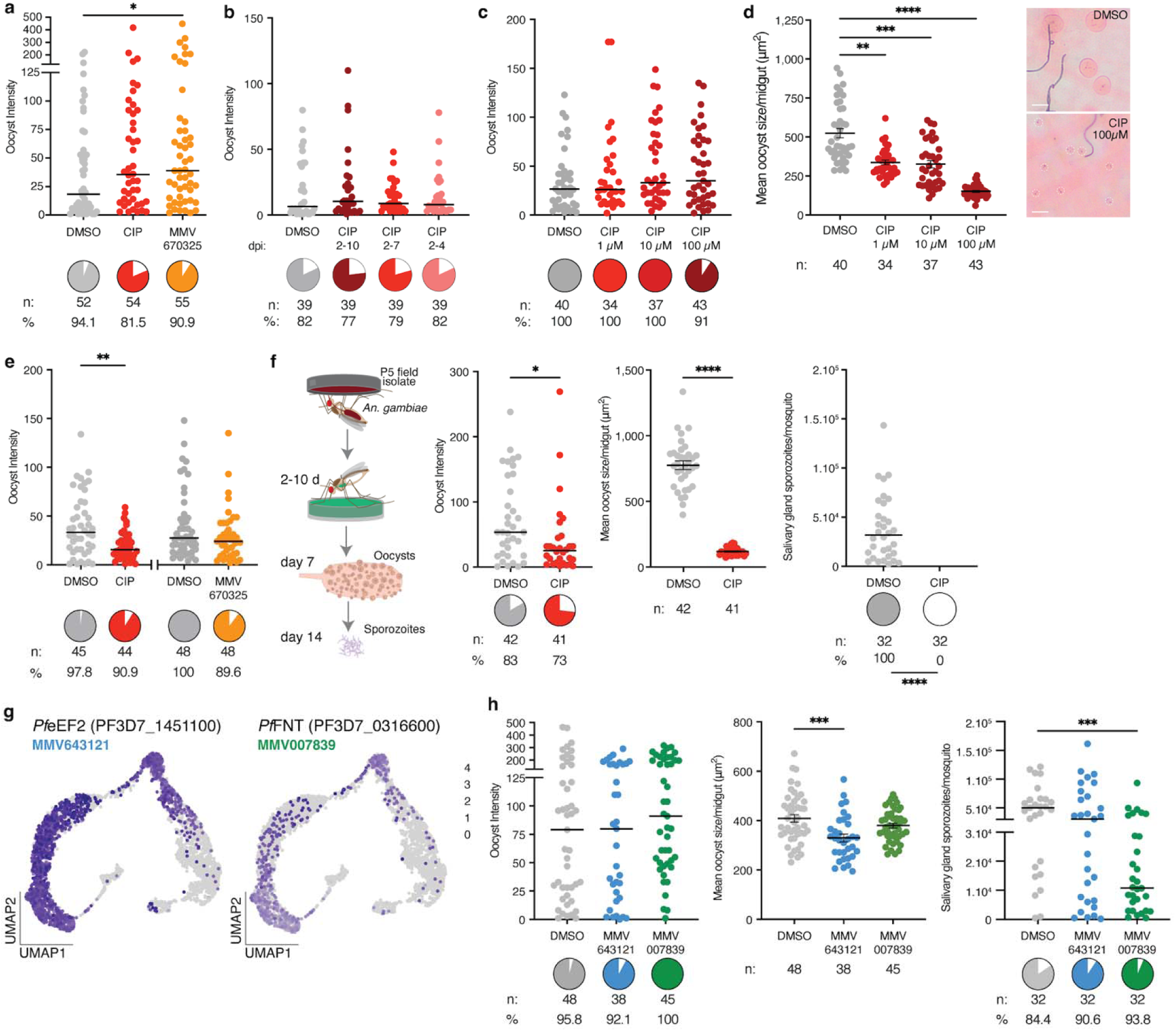
Functional validation of candidate genes in oocyst growth and sporozoite infection. **a,** CIP and MMV670325 delivered in sugar from 2-10d pi to *An. gambiae* infected with NF54 had no effect on oocyst prevalence, while MMV670325 treatment slightly increased oocyst intensity (p=0.041). **b-d,** CIP exposure for three different **(b)** durations or **(c-d)** concentrations had no effect on **(b-c)** oocyst prevalence or intensity but **(d)** significantly reduced oocyst size in a dose dependent manner (p=0.0033, p=0.0002, p<0.0001; representative images of DMSO and 100 µM CIP treatment in right panel, Scale bar: 20 µm). **e**, The same treatments to *An. stephensi* infected with the ART29 isolate resulted in no prevalence difference, but a slight decrease in oocyst intensity (p=0.002) after CIP treatment. **f**, CIP treatment in *An. gambiae* infected with the P5 isolate slightly decreased oocyst numbers (p=0.016), but significantly reduced oocyst size (p<0.0001) at 7d pi, resulting in no sporozoites in the salivary glands at 14d pi. **g**, Expression profile of *Pf*eEF2 and *Pf*FNT, targets of MMV643121 and MMV007839, respectively. **h**, Exposure to either drug did not affect oocyst prevalence or intensity. MMV643121 treatment led to a decrease in oocyst size (p=0.0005), while MMV007839 did not affect oocyst size but resulted in reduced sporozoite intensity (p=0.0009). Oocyst size data are represented as mean ± SEM, oocyst and sporozoite intensity data as medians. All are compared using either a Mann Whitney two-tailed test (panels e and f) or a Kruskal-Wallis test and Dunn’s correction (all other panels). *n* indicates the number of mosquitoes from at least two independent infections. Pie charts indicate infection prevalences, compared with two-tailed Fisher’s exact test. *p<0.05, **p<0.01, ***p<0.001, ****p<0.0001. Schematics in **f** are adapted from ref. 36 under a Creative Commons licence CC BY 4.0.

**Extended Data Figure 5.**
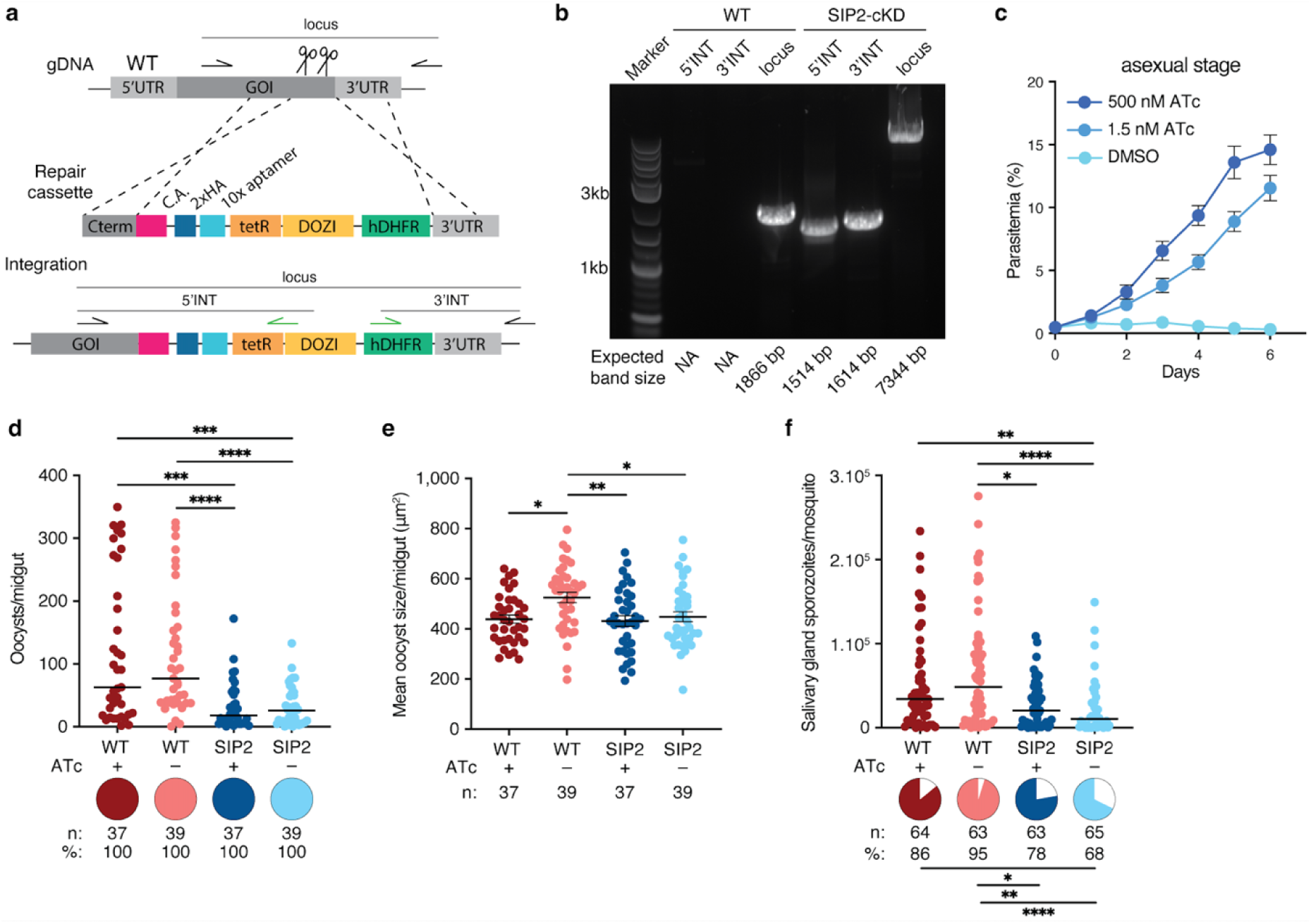
Conditional knockdown of PfSIP2 does not affect sporozoite intensity. **a**, Scheme (not to scale) of the *Pf*SIP2 conditional knockdown (cKD) construct. Positions of primer pairs for integr tion validation are indicated. **b,** PCR amplifications of the 5’ and 3’ integrated regions (5’INT and 3’INT) and locus in WT NF54 and *Pf*SIP2-cKD parasites, demonstrating the absence of WT parasites in the cKD line. **c,** *Pf*SIP2 knockdown by ATc withdrawal in the asexual blood stage led to parasite death (Mixed-effects model, Dunn’s correction, p<0.0001). **d-f,** PfSIP2 knockdown by ATc withdrawal had no effect on (**d**) oocyst prevalence (Fisher’s exact test, two-tailed), intensity (Kruskal-Wallis test, Dunn’s correction, p=0.0004, p=0.0007, and p<0.0001), and (**e**) size (Kruskal-Wallis test, Dunn’s correction, p=0.0152, p=0.0082, and p=0.0258) at 7d pi, and had no effect on (**f**) prevalence (p=0.0211, p=0.0076, and p<0.0001) and intensity (p=0.0024, p=0.0213, and p<0.0001) of salivary gland sporozoites at 14d pi (two-tailed Fisher’s exact tests and Kruskal-Wallis test with Dunn’s correction, respectively). Data are represented as mean ± SEM in panels **c,** and **e**, and as median in **d,** and **f**. *n* indicates the number of mosquitoes and pie charts and % infection prevalence. *p<0.05, **p<0.01, ***p<0.001, ****p<0.001.

**Extended Data Figure 6.**
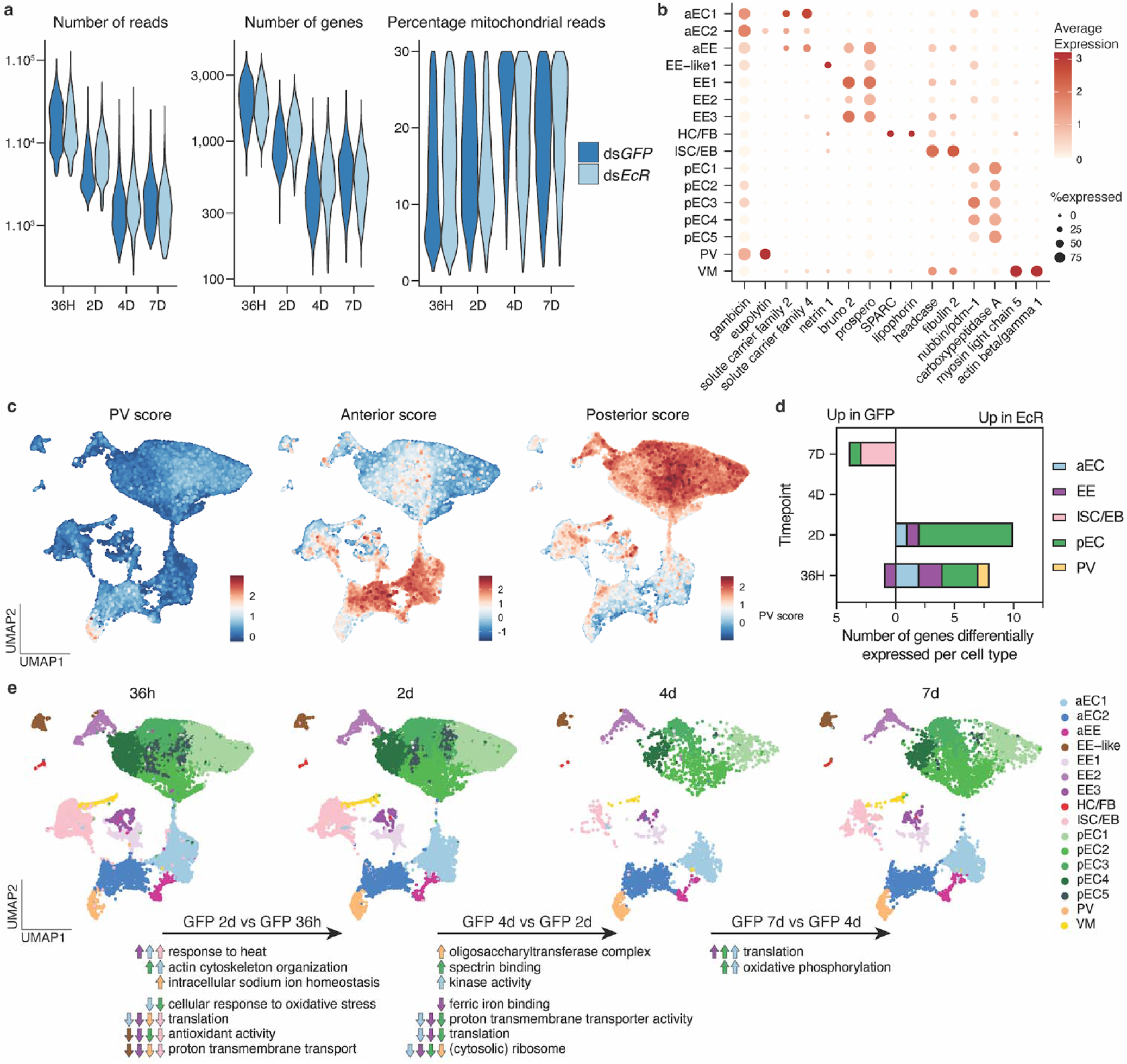
Quality control and clustering of the *An. gambiae* scRNA-seq data. **a**, Violin plots displaying key quality metrics for the mosquito scRNA-seq datasets (left to right: total number of reads, of genes, and percentage of mitochondrial reads) after QC. **b**, Dot plot displays cell type-specific markers, showing their average expression (colour) and the proportion of expressing each marker (size). Cell types shown are anterior enterocytes (aEC), enteroendocrine (EE), EE-like cluster, hemocytes and fat body (HC/FB), intestinal stem cells/enteroblasts, (ISC/EB), posterior enterocytes (pEC), proventriculus or cardia (PV) and visceral muscles (VM). **c**, Expression of markers characteristic to the proventriculus, and anterior or posterior part of the midgut based on study by Hixson *et al.*^51^. **d**, Number of genes differentially expressed by pseudobulk analysis between ds*GFP* and ds*EcR* treated mosquitoes per time point and cell type (Wald test, significance defined as adjusted p-value < 0.05 and absolute value of log_2_ fold change > 0.5). **e**, UMAPs of *An. gambiae* cells split by time point, showing the dynamic changes of the midgut cell population shortly after blood digestion (36h and 2d pi) and when subsequently maintained on sugar-only diet (4d and 7d pi). Pseudobulk differential expression analysis was performed in ds*GFP-*treated mosquitoes by pooling each cell type and comparing consecutive time points. Changes to significant GO term enrichment are represented by up and down arrows, colored by cell type.

**Extended Data Figure 7.**
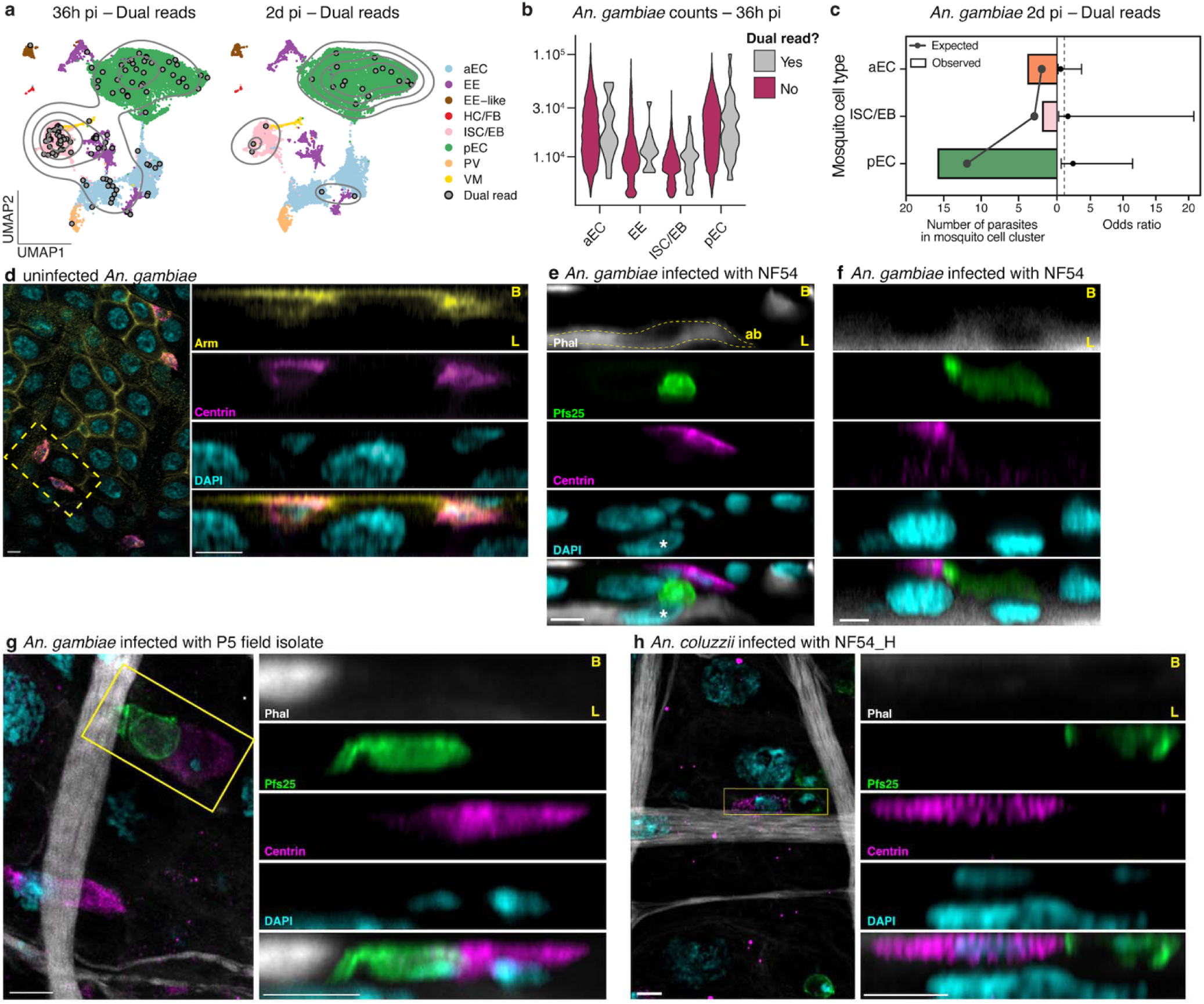
Dual-read parasites at 36h pi show an association with intestinal progenitor cells. **a,** *An. gambiae* UMAPs at 36h and 2d pi colored by cell type and overlaid with dual-read cells (gray circles) and contour lines (light gray lines) displaying the density of dual-read cells. **b,** Violin plot showing similar read counts in mosquito cells regardless of whether they are dual-read cells. **c,** twenty-one dual-read parasites over eight independent samples at 2d pi were randomly distributed across mosquito cell types, with no differences in (left x-axis) the observed vs. expected numbers and (right x-axis) resulting odds ratio (point) ± 95% confidence interval (error bar). Fisher’s exact test, two-tailed, with BH multiple comparison correction. **d,** Maximum intensity projection from confocal imaging (9.66 µm Z-stack) showing colocalization of the centrin (magenta) signal with the well-characterized ISC marker, armadillo (Arm, yellow), located on the basal side of the midgut (B, top). **e-f,** Confocal imaging of NF54 parasites (Pfs25, green) traversing the *An. gambiae* midgut epithelium at 36h pi. (**e**) Parasites were found to point towards a progenitor cell (centrin, magenta) directly above an extruding cell (asterisk, nucleus of the extruded cell), located between two actin brushes (ab, yellow dashed lines) delimiting each epithelium (phalloidin, gray, gamma 0.5). (**f**) Ookinete left a Pfs25 trail into a large cell reminiscent of an enterocyte (see Supplementary Video 1). **g-h,** The interaction was also observed in (**g**) *An. gambiae* midguts infected with P5 parasites and (**h**) *An. coluzzii* midguts infected with NF54_H parasites (max intensity projection of 1.69 and 1.68 µm Z-stacks, respectively). Imaging was performed on at least two independent infections. Phalloidin (phal, gray) was used to define the basal side (B, top) and the luminal brush border (L, bottom), and nuclei were stained with DAPI (cyan). Scale bar: 5 µm.

**Extended Data Figure 8.**
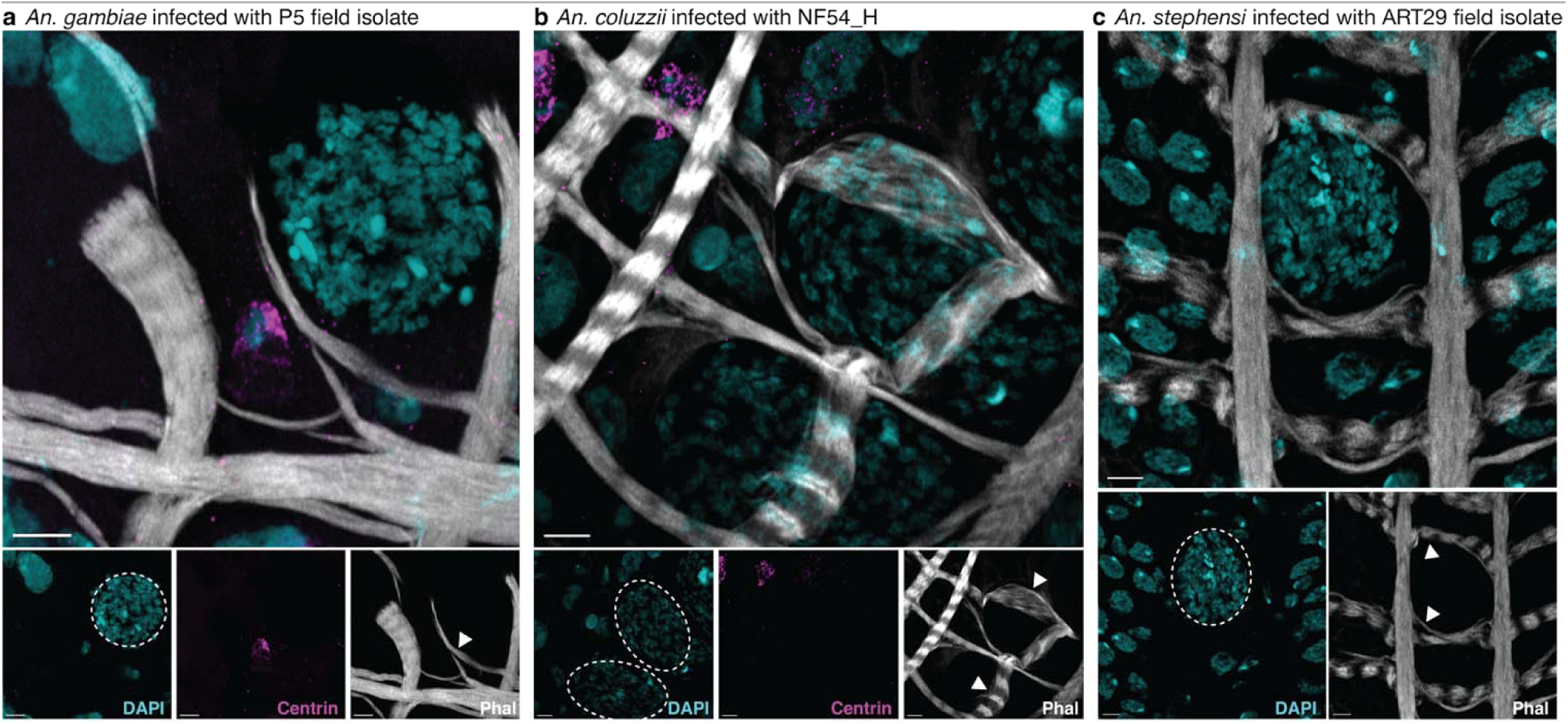
Visceral muscle wraps around late-stage oocysts. **a-b**, Visceral muscle (Phal, gray) wrapped closely around late oocysts (arrowheads), with no apparent increase in progenitor cell (centrin, magenta) populations near oocysts in (**a**) *An. gambiae* midguts infected with P5 parasites and (**b**) *An. coluzzii* midguts infected with NF54_H parasites by confocal imaging (max intensity projection of 4.60 µm and 2.60 µm Z-stacks, respectively). **c**, Similar muscle deformation was observed in *An. stephensi* midguts infected with ART29 parasites (max intensity projection of 3.60 µm Z-stack), though inconsistent centrin staining prevented analysis of progenitor cell distribution. At least two independent infections were used for imaging. Scale bar: 5 µm.

## Supplementary Note

### Parasite sequencing depth requirements

Each replicate of the eight samples was mixed equally and initially sequenced on an Illumina iSeq 100 to determine the percentage of reads mapping to parasites relative to mosquito cells. As stated in the methods, around 0.05%, 0.1%, 0.6%, and 1.3% reads mapped to parasites at 36h, 2d, 4d and 7d pi, respectively. Based on hemocytometer counts, we estimated that each sample loaded onto the 10X chip contained an average of 380 parasites, resulting in an estimated total of 3,040 parasites per time point (from two metabolic conditions and four replicates). Based on these estimates, we selected two NovaSeq S4 flow cells, which were expected to yield a minimum of 12 billion reads (Broad Institute). This would provide an estimated 6-156 million reads mapping to parasites at 36h to 7d pi, corresponding to an estimated reading depth of 1,974–51,316 reads per parasite at 36h to 7d pi.

### Canonical midgut cell types and functional validation in hematophagous mosquitoes

The mosquito midgut is divided into anterior and posterior regions, with the latter making up the bulk of the organ and serving as the site for blood meal storage and digestion^1^. Situated at the anterior-most end of the midgut, the proventriculus (PV), also known as the cardia, is a distinct structure involved in immune responses in adult mosquitoes^2,3^. Data on midgut cell types in hematophagous mosquitoes is relatively sparse (reviewed by Hixson *et al.*^4^) but is extensively available from studies of *Drosophila melanogaster*^5–8^, and has been used to annotate cell types of other mosquito midgut scRNA publications. Canonically, these include intestinal stem cells (ISCs), undifferentiated enteroblasts (EBs), and the terminally differentiated enterocytes (ECs), enteroendocrine cells (EEs), and visceral muscle cells (VMs).

ISCs are the only replicative cells in the midgut epithelium, underpinning both tissue homeostasis and regenerative capacity. Located basally, with no direct access to the gut lumen, ISCs undergo either symmetrical—yielding two self-renewing ISCs—or asymmetrical division, producing one ISC and one differentiating daughter cell^9–11^. The existence of such stem cells in mosquito midguts was long debated but active mitosis has since been imaged in the midguts of *Anopheles* mosquitoes^12–14^. A recent study extensively characterized external stimuli, such as a blood meal or bacterial infection, which trigger ISC division in *An. gambiae* and *Ae. aegypti*^14^.

Following asymmetric division, the daughter cell fated for differentiation may adopt one of two trajectories: it can become an EB—an intermediate, undifferentiated cell that will subsequently differentiate into an EC^11,12^—or a pre-EE that will mature into a functional EE^15^. Although EEs have been described in midgut images for decades^16–18^, recognized by their abundant vesicles and basolateral exocytosis, their functional roles remain poorly characterized. A subset of these cells expresses neuropeptide F, a hormone implicated in stimulating feeding behaviour in *Ae. aegypti*^19^, suggesting that EEs may serve as key regulators of host attraction and gut-brain communication.

ECs constitute the majority of the epithelium in both the anterior and posterior midgut. These large, columnar cells exhibit microvilli on their luminal surface, supporting their roles in digestive enzyme secretion and nutrient absorption^12,16,20^. ECs are polyploid, a feature particularly accentuated in *An. gambiae*, where nuclear content can reach up to 256n following a blood meal^14^. Their function varies according to their location, with anterior ECs being responsible for sugar digestion and absorption while the posterior ECs mainly produce enzymes for blood meal digestion and uptake amino acids^1,3,20^.

Although not epithelial in origin, VMs are essential for midgut integrity. A lattice of longitudinal and circular muscle fibers envelops the basal surface of the midgut epithelium^21^, enabling coordinated peristalsis and contributing to the mechanical plasticity of the organ during blood meal intake and digestion.

### Mosquito scRNA cell cluster annotation

Mosquito single-cell data were annotated using a combination of single-cell (sc) and single-nucleus (sn) midgut studies, alongside bulk RNA-seq data from *Anopheles* and functional studies primarily conducted in *D. melanogaster* (reviewed by Zhang and Edgar^8^). First, cell localization was scored based on bulk RNA-seq expression profiles of distinct regions of the digestive tract, which includes the proventriculus (PV), and the anterior and posterior midguts^1^. This analysis led to the identification of cluster 10 as PV (Extended Data Fig. 6c).

We then compared our cluster markers with those reported in sn or scRNA-seq studies of *D. melanogaster*, *Ae. aegypti* and *An. gambiae*^5,6,22,23^. Given the comprehensiveness of the Fly Cell Atlas, which encompasses all *D. melanogaster* cell types, we limited the reference dataset to midgut-relevant populations: ISCs, EBs, VMs, EEs, ECs (anterior and posterior), cardia or PV, along with likely contaminants such as hemocytes (HC), fat body (FB), Malpighian tubule cells, and tracheal cells. By overlapping the expression of reference marker genes with our cluster marker genes (Supplementary Table 5), we observed a robust signal for EE in mosquito cell clusters 8, 9, 12, and 13; ISC/EB in cluster 4; and VM in cluster 14. Signal from HC and FB remained mainly confined to the smallest cluster 15, consistent with their status as contaminants. Markers for Malpighian tubules, cardia, PV, and tracheal cells were largely absent, including in the previously annotated PV cluster. The remaining clusters were presumed ECs as they express EC-associated markers, even though expression of these markers was broadly distributed across multiple clusters.

Functional enrichment of each cluster was next used to validate cluster annotation (Supplementary Table 5). All EE-annotated clusters showed enrichment in biological processes related to cell communication and cell–cell signaling, consistent with their endocrine function. The ISC/EB cluster was enriched for the terms ‘mitotic cell cycle’ and ‘DNA replication’, reflecting its proliferative capacity, while the VM cluster showed enrichment for genes involved in sarcomere organization and muscle contraction. The PV cluster expressed high levels of genes associated with humoral immune response, in agreement with previous studies and its anticipated role in anti-microbial peptide production^1–3,6^. As expected for a cluster comprising two distinct cell types, the functional enrichment profile of the HC/FB cluster was less informative. Clusters 0, 1, 3, and 7, which showed high posterior scores, were enriched in proteolysis and metalloaminopeptidase activity, consistent with pEC identity. Clusters 2 and 5, both with high anterior scores, were enriched for heme-binding proteins—hallmarks of aECs^1^—and were annotated accordingly. Interestingly, cluster 6 showed a high posterior score but was enriched in sugar transporters, contrary to the notion that such function is restricted to the anterior midgut. This may reflect a greater investment of *An. gambiae* on carbohydrate digestion and absorption in the posterior midgut compared to *Ae. aegypti*, and the cluster was annotated as pEC^1^. Finally, cluster 11 was enriched for genes involved in amide and sulfur compound metabolism, but these functions have not been clearly linked to a specific midgut cell type.

Finally, we validated cluster annotations with established markers reported in the literature (Extended Data Fig. 6b). Delta and klumpfuss (klu)—canonical markers of ISCs and EBs, respectively^11,24^—were only weakly detected in our dataset, though headcase^25^, a known ISC marker, emerged as one of the top five markers in the ISC cluster, supporting its annotation. The VM cluster showed high expression of myosin and actin genes, while the HC/FB cluster expressed SPARC and lipophorin, consistent with their known marker profiles^22^. All pEC clusters showed high levels of nubbin^5,8,23,26^ and carboxypeptidase A, with low levels of gambicin. In contrast, aEC and PV clusters exhibited strong expression of gambicin; aECs also expressed sugar transporters and digestive enzymes, while PV was marked by a distinct eupolytin. EE clusters expressed prospero, a well-established EE marker in *D. melanogaster* and *Ae. aegypti*^5,23,26^, as well as bruno 2, identified as an EE marker in the latter species. Cluster 11 expressed prospero but lacked expression of most other EE markers, including bruno 2, and was therefore annotated as EE-like.

Together, this single-cell transcriptomic analysis enabled systematic identification and characterization of midgut cell types. For the first time in mosquito midgut studies, spatial positioning of individual cells and clusters was inferred from empirical data^1^, revealing an anterior EE cluster previously identified through imaging^17,18^.

## Supplementary Tables

Supplementary Table 1: Percentage of reads mapped to the genome of *An. gambiae* and *P. falciparum*, and quality control criteria for each sample.

Supplementary Table 2: Pseudobulk gene expression and functional enrichment of *P. falciparum*.

Supplementary Table 3: Marker genes for each cluster of the integrated *P. falciparum* midgut stages dataset.

Supplementary Table 4: Gene model of *P. falciparum* midgut stages from this study.

Supplementary Table 5: Top 100 marker genes and corresponding functional enrichment of each cluster of the integrated *An. gambiae* midgut cells and proportion across timepoints.

Supplementary Table 6: Pseudobulk differential expression analysis comparing mosquito cells in ds*EcR* and ds*GFP* treatment at each time point.

Supplementary Table 7: Pseudobulk gene expression and functional enrichment of *An. gambiae* midgut cells across time points post blood meal.

Supplementary Table 8: Primer information.

## Supplementary Videos

Supplementary Video 1: 3D rendering of the Extended Data Figure 7f illustrating an ookinete that has left a Pfs25 trail into a large cell reminiscent of an enterocyte. DNA was labelled with DAPI (cyan), VM with phalloidin (grey), ISC with centrin (magenta) and parasite with Pfs25 (green).

Supplementary Video 2: Basal to lumen video of successive slices from the Fig 5a Z-stack illustrating the muscle stretch around and oocyst. DNA was labelled with DAPI (cyan), VM with phalloidin (grey), and ISC with centrin (magenta). Scale bar: 10 µm.

Supplementary Video 3: Lumen to basal video of successive slices from the Fig 5c Z-stack illustrating the muscle lattice encasing an oocyst. DNA was labelled with DAPI (cyan), and VM with phalloidin (grey).

